# Tripartite separation of glomerular cell-types and proteomes from reporter-free mice

**DOI:** 10.1101/2020.08.22.262584

**Authors:** Favian A. Hatje, Markus M. Rinschen, Uta Wedekind, Wiebke Sachs, Julia Reichelt, Tobias B. Huber, Sinah Skuza, Marlies Sachs, Stephanie Zielinski, Catherine Meyer-Schwesinger

**Affiliations:** Institute of Cellular and Integrative Physiology, University Medical Center Hamburg-Eppendorf, 20246 Hamburg, Germany; III Department of Medicine, University Medical Center Hamburg-Eppendorf, 20246 Hamburg, Germany; Department of Biomedicine, Aarhus University, Aarhus, Denmark; Department II of Internal Medicine, and Center for Molecular Medicine Cologne, University of Cologne, Cologne, Germany

**Keywords:** Glomerulus, podocyte, mesangial cell, glomerular endothelial cell, TimMEP, isolation, species, artifact

## Abstract

**Purpose:** The kidney glomerulus comprises a syncytium of podocytes, mesangial and endothelial cells, which jointly determine glomerular filtration barrier function, and thereby kidney and cardiovascular health. The understanding of this intricate functional unit and its intracellular communication beyond the transcriptome requires bulk isolation of these cell-types from glomeruli for subsequent biochemical investigations. Therefore, we developed a globally applicable tripartite isolation method for <u>m</u>urine <u>m</u>esangial and <u>e</u>ndothelial cells and <u>p</u>odocytes (timMEP).

**Methods:** Glomerular cells were separated via a novel FACS-sort depending on a cell-specific antibody labeling in wildtype mice or based on a combination of transgenic fluorescent protein expression and antibody labeling in mT/mG mice. The purity of isolated cell-types was validated by qPCR and immunoblot. The proteome of podocytes, mesangial and endothelial cells was determined and compared between species, ages and gender of wildtype and mT/mG mice. The method was also applied to the podocyte-targeting immunologic injury model of THSD7A-associated membranous glomerulonephritis.

**Results:** TimMEP enabled protein-biochemical analyses of podocytes, mesangial and endothelial cells derived from a single reporter free mouse. Proteomic analyses allowed the first characterization of podocyte, endothelial and mesangial proteomes of individual mice. Marker proteins for mesangial and endothelial proteins were determined, and protein-based interaction and intraglomerular cell communication networks were elucidated. Interestingly, analyses revealed significant cell-type specific proteome differences between mouse strains, artefacts induced by reporters, and alterations depending on gender and age. Within the glomerulus, timMEP resolved a fine-tuned initial stress response exclusively in podocytes after exposure to anti-THSD7A antibodies, which was not detectable using conventional analyses in whole glomeruli.

**Conclusion:** Globally applicable timMEP abolishes the need for costly, time- and animal-consuming breeding of mice to glomerular cell-type reporters. TimMEP enables glomerular cell-type resolved investigations at the transcriptional and protein biochemical level in health and disease, while avoiding reporter-based artefacts, paving the way towards the comprehensive and systematic characterization of glomerular cell-type biology.

**Key messages:**

1. A tripartite isolation method for mesangial, endothelial and podocyte cell-types from reporter-free mice.
2. Generation of bulk cell-type samples and primary co-cultures for biochemical and protein-based analyses.
3. Strain and transgene-dependent expression of proteins among glomerular cell-types, including protein profiles, intra-glomerular communication machineries, and reporter-dependent artefacts.
4. Disease specific time-resolved resolution of glomerular cell-type’s response to injury.

## INTRODUCTION

The glomerular filtration barrier represents a sophisticated syncytium of individual cell types, namely podocytes, mesangial and glomerular endothelial cells, which sustain the structure and regulate the function of the filtration barrier (Pavenstadt et al., 2003). Podocytes embrace the glomerular capillaries and form an intricate mesh with their interdigitating processes that are interconnected by a modified form of adherens junction, the slit diaphragm, which ultimately bridges the filtration slits. The intraglomerular mesangial cells are situated in close contact with the endothelial cells and represent a specialized form of pericytes that provide structural support. The fenestrated endothelial cells line the glomerular capillaries and reside opposite from the podocytes separated by the glomerular basement membrane. Malfunction of any of these cell types leads to loss of glomerular function and proteinuria leading to the concept that these glomerular cell types interact (Dimke et al., 2015), a finding also supported in silico by single-cell transcriptomics studies. However, single cell approaches are still mRNA based, and limitations include lack of depth of transcriptomes, and the difficulty to subject single cells towards biochemical applications. Understanding glomerular crosstalk would require the bulk isolation of all three cells-types from glomeruli of individual mice for subsequent protein biochemical investigations.

The establishment of a globally applicable large scale glomerular isolation method from mice using magnetic beads (Takemoto et al., 2002) has revolutionized our understanding of glomerular injury pathways. Later, a FACS-based isolation method of podocytes from mice (Wanner et al., 2014) has further enhanced the cellular resolution of glomerular injury patterns with an isolated focus on podocytes. This technique relies on the use of mT/mG mice, which exhibit a transgenic intracellular expression of eGFP in podocytes and of tdTomato in non-podocytes under control of a Cre promotor (Muzumdar et al., 2007). However, this method has significant limitations. First, prior to podocyte isolation, mouse models of interest are required to be crossed to mT/mG mice, resulting in a costly and time- and mouse-consuming breeding, with the risk of genetic background changes, which are often accompanied by a change in susceptibility to established injury models. Further, only podocytes are isolated from one mouse, the other glomerular cells being indistinguishable through the expression of tdTomato. Additionally, the substantial expression of a foreign fluorescent protein within glomerular cells (up to 6%) (Rinschen et al., 2018a) could induce inevitable biochemical artefacts, depending on the biological system evaluated (for example protein degradation systems). These limitations are particularly important in diseases with altered glomerular composition such as podocyte loss or mesangial expansion.

The aim of this study was to develop a 1) globally applicable and 2) cost / time / mouse economizing isolation protocol of all three glomerular cell types from one individual mouse in sufficient amounts and purity to enable physiological and pathophysiological protein biochemical and omics investigations of *in vivo* mouse models.

## MATERIALS AND METHODS

### Animals

Male and female C57BL/6 mice and BALB/c mice were purchased from Charles River (Germany) at the age of 10-14 weeks. mT/mG mice (ICR;Sv129/J;C57BL/6) (Boerries et al., 2013) were provided from the Huber laboratory (III Medical Clinic, UKE, Hamburg-Eppendorf). Mice had free access to water and standard chow, were synchronized to a 12:12 hour light-dark cycle and sacrificed by cervical neck dissection for kidney removal. Anti-THSD7A membranous nephropathy was induced by intravenous injection of 180 µl rabbit anti-THSD7A antibodies or unspecific rabbit IgG as control in male BALB/c mice aged 12-22 weeks (Tomas et al., 2017). Urine was collected as spontaneous urine on day 1 and day 7 prior to sacrifice.

### Measurement of proteinuria

Urine samples were collected over 3-5 hours in a metabolic cage with free access to water. Urine albumin content was quantified using a commercially available ELISA system (Bethyl) according to the manufacturer’s instructions, using an ELISA plate reader (BioTek), as described (Meyer et al., 2007). Values were standardized against urine creatinine values of the same individuals determined according to Jaffe and plotted.

### Glomerular cell isolation

Kidneys were harvested, perfused with magnetic DynaBeads and glomeruli were isolated as described before (Takemoto et al., 2002). Collected and pelleted glomeruli were dissolved in digestion buffer. The digestion buffer contains 1000 µg/ml Liberase TL (Roche), 100 U/ml DNase1 (Roche), 10% FCS, 1% ITS, 1% PenStrep, 25 mM HEPES dissolved in RPMI Medium 1640 1x (Gibco). Cells were incubated for 2 hours at 37°C, 1400 rpm and repeatedly and diversely mechanically stressed by vortexing, shearing (with a 27G needle) and pipetting using Pasteur pipettes and Eppendorf pipettes to promote maximum cellular separation. A DynaMag was used to separate glomerular remnants and DynaBeads from singular cells. After 5 minutes in the DynaMag, the single cell containing supernatant was collected. Cells were pelleted (10 min, 4°C, 1000 g) and washed once with MACS (PBS with 0,5% BSA and 2 mM EDTA). Following centrifugations, the supernatant was removed as carefully as possible to avoid any disturbance of the very fragile cell pellet. Subsequently, cells were stained with cell-specific antibodies podoplanin (podocytes), CD73 (mesangial cells), CD31 (endothelial cells) as well as CD45 (leukocytes) and a live/dead stain (Invitrogen) listed in ***Supplemental table 1***. Following the incubation (30 minutes at 4°C in complete darkness) cells were washed once more, resuspended in MACS and sieved through 40 µm sieves into the FACS tubes. Cells for the proteomics experiments in this paper were then sorted with the strategy shown in Supplement Figure 10A. However, since then we revised the strategy to compensate for slight cross-contamination of podocytes in the mesangial cells, (***Supplemental figure 1B, C***) and we only recommend using the strategy shown in Figure 1D for sorting.

**Figure 1:**
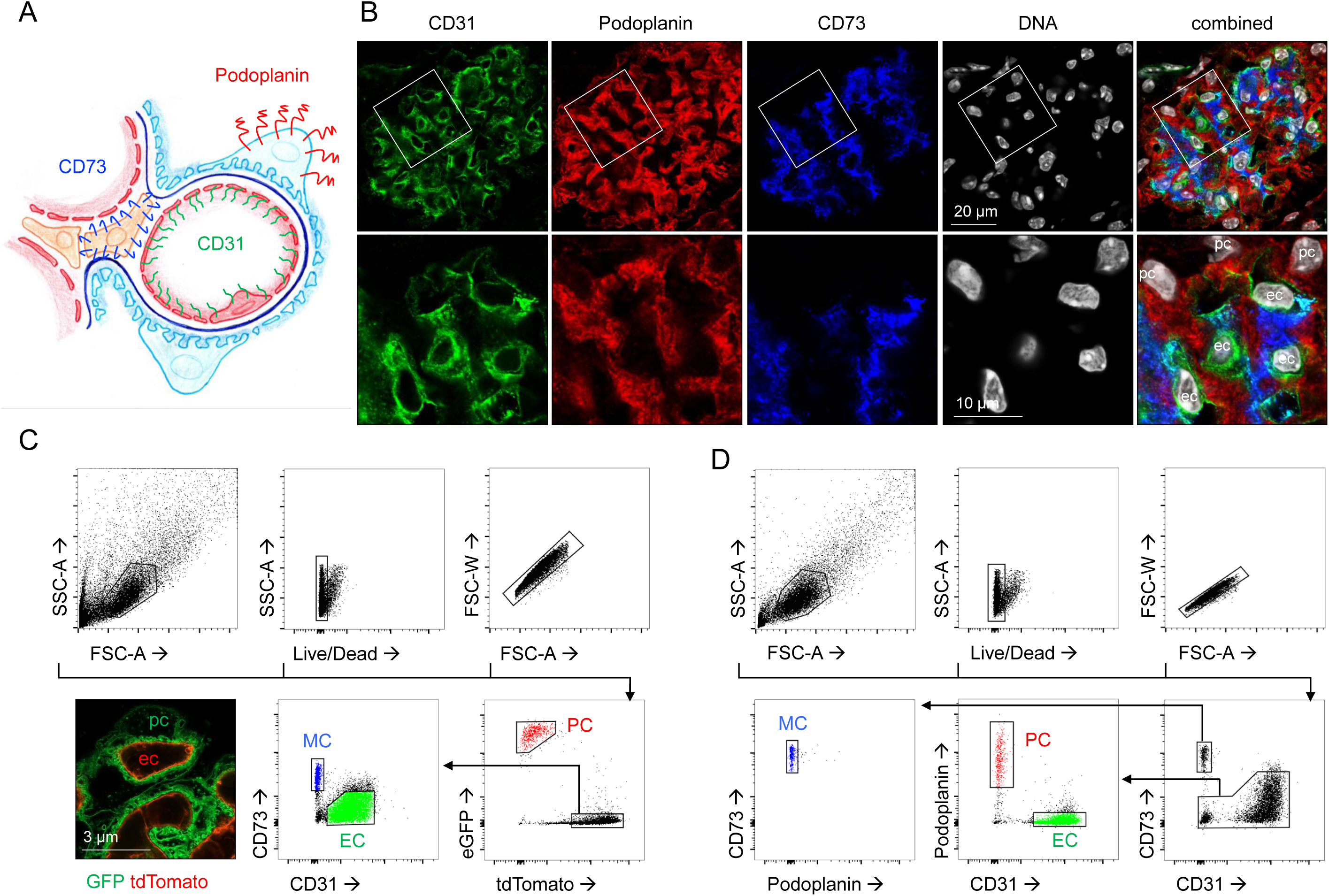
FACS-sorting strategy from wildtype mice and mT/mG mice for tripartite isolation of glomerular cell types. (**A**) Scheme of a glomerular loop depicting the localization of the cell specific markers used for the FACS-sorting strategy. (**B**) Confocal micrographs depicting labeled podocytes (podoplanin, red), endothelial cells (CD31, green) and mesangial cells (CD73, blue) in a 4% PFA fixed frozen section of a murine kidney, DNA was visualized using Hoechst, pc = podocyte, mc = mesangial cell, ec = endothelial cell. (**C**) Gating strategy of glomerular single cells derived from a mT/mG mouse demonstrating a distinct podocyte (red; eGFP^+^/tdTomato^−^), endothelial cell (green; tdTomato^+^/eGFP^−^ /CD31^+^/CD73^−^) and mesangial cell (blue; tdTomato^+^/eGFP^−^/CD73^+^/CD31^−^) population. The unlabeled cells represent contaminating cells such as tubular cells. The left lower panel at the lower left exhibits a representative high-resolution confocal image of the intrinsic GFP and tdTomato expression in a mT/mG glomerulus from an optical cleared kidney section. The histological panel exhibits intrinsic GFP and tdTomato fluorescence of a glomerular capillary loop of an optically cleared 300 µm kidney slice. (**D**) Gating strategy of glomerular single cells isolated from a wildtype mouse exhibiting a distinct podocyte (PC, red; podoplanin^+^/CD73^−^/CD31^−^), endothelial cell (EC, green; CD31^+^/CD73^−^/podoplanin^−^), and mesangial cell (MC, blue; CD73^+^/CD31^−^/podoplanin^−^) population. The unlabeled cells represent contaminating cells such as tubular cells.

### Sample preparation and mass spectrometry analysis

Cell pellets were snap-frozen and stored at −80 degrees. For comparison of different strains, approximately 50000 cells were analyzed. Cells were resuspended in 4% SDS and 10mM Tris and heated at 95 degrees for 10 min. Then, solubilized proteins were reduced and alkylated using 5mM DTT and 10mM Iodoacetate, respectively. Proteins were digested and prepared using the SP3 protocol with modifications previously described (Hohne et al., 2018; Rinschen, 2019). For deeper protein analysis, pellets of 1 million cells were obtained from mT/mG mice. These pellets were resuspended in 8M urea and 5mM Tris in LCMS grade water, sonicated for 1 min using an ultrasonication pulse (0.1 sec cycle, 10% strength), and spun down with 16000*g* at 4°C. Protein concentration (for 1e6 cells) was determined using a commercial BCA assay (Thermo). Then proteins were reduced and alkylated as described above. Proteins were digested using a protease in-solution digestion protocol with a modified SP3 protocol (Hohne et al., 2018; Hughes et al., 2019) (50000 cells), or in solution digestion (Rinschen et al., 2018a). For all digestion steps, trypsin was used. Tryptic peptides were analyzed using a nLC-MS/MS hardware setup consisting of a nanoflow LC (flow: 200 nl/min) coupled to an Orbitrap QExactive Plus tandem mass spectrometer. The peptides were separated using a gradient used for reverse phase separation consisted of buffer A and buffer B, with ascending concentrations of buffer B (80% acetonitrile, 0.1% formic acid) over buffer A (0.1% formic acid). The peptides from 50000 cells were separated using a 1h gradient. The peptides from 1*e6 cells were separated using a 2.5h gradient.

### Bioinformatic analysis

Protein raw files were searched using MaxQuant and the LFQ algorithm (Cox et al., 2014; Cox and Mann, 2008) with search against a uniport mouse proteome reference database released in Jan 2018. Search criterions were alkylation on cysteins as a fixed modification, and N-terminal acetylation as well as methionine oxidation as variable modifications. Default criterions were used, meaning that PSM, peptide and protein FDRs were set at 0.01. LFQ algorithm was enabled, match between run was enabled. The data were analyzed using Perseus v 1.5.5.3 using filtering for the embedded annotations as contaminant, reverse or proteins identified by site only. Only proteins in at least 60% of samples were kept and missing values were imputed using default imputation parameters (downshift SD=2, width 0.3). GO-term and uniport keyword annotation and enrichment was performed using the embedded terms (Tyanova et al., 2016). Radarplots were generated using ggradar package (Rstudio) https://github.com/ricardo-bion/ggradar using default settings. Generated txt files are shared (raw data deposition). Differentially proteins were defined by ANOVA with FDR-corrected *p* value of less than 0.01 to adjust for multiple testing. Among these proteins, criteria defined for the identification of new glomerular cell-type enriched proteins: 1) a Student t-test difference cut-off of > 2 between for example podocytes and non-podocytes, 2) a negative search result in pubmed (https://www.ncbi.nlm.nih.gov/pubmed) with the keywords “protein of interest” and “podocyte” or “glomerular” or “renal” or “kidney”; “protein of interest” and “mesangium” or “mesangial” or “glomerular” or “renal” or “kidney”; and “protein of interest” and “endothelial” or “endothelium” or “glomerular” or “renal” or “kidney” 3) a validated cell of interest expression pattern in the human protein atlas (https://www.proteinatlas.org/search). From significantly enriched podocytes, protein-protein interaction networks were generated using STRING db (Szklarczyk et al., 2017), and filtered for high confidence (more than 0.9) based on experimental or database evidence only. Then, protein networks were imported into cytoscape v 3.3 (Shannon et al., 2003), and all edges that were not between two different cell types and the associated nodes were removed.

### Raw data deposition

The raw data of the files is available at Pride/ProteomExchange at the link https://www.ebi.ac.uk/pride/archive/login using the following tokens (Perez-Riverol et al., 2019; Vizcaino et al., 2016)

Dataset: mT/mG dataset

**Project Name:** Analysis of mouse glomerular cells

**Project accession:** PXD016238

**Project DOI:** Not applicable

**Password:** UHUYumfS

Dataset: Different strains

**Project Name:** FACS-sorted podocytes, mesangial cells

**Project accession:** PXD016237

**Project DOI:** Not applicable

**Password:** kQUh8pjC

### qPCR analysis

Total messenger RNA was extracted from FACS-sorted cells and isolated glomeruli using NucleoSpin RNAII kit and NucleoSpin RNA Plus XS (both Machery-Nagel) according to the manufacturer’s instructions and was reverse transcribed with random hexamer primer (Invitrogen) and Revert Aid (Thermo Fisher). mRNA expression was quantified with QuantStudio 3 using SYBR green as recently described (Panzer et al., 2006). Exon spanning primer pairs to murine cDNA are listed in ***Supplemental table 2***. 18S was used as an internal control to correct for small variations in RNA quality and cDNA synthesis essentially as described by AbiPrism. Amplicons of random samples for each primer pair were determined by automatic PCR sequencing to demonstrate the specificity of the PCR reaction (data not shown). ΔCT values were calculated using 18S as homekeeper. ΔΔCT values were calculated as difference between glomerular and sorted cell ΔCT, RE (relative expression) is 2^(ΔΔCT). Displayed are RE of sorted cells in percent of the glomerular RE, calculated as follows: 100% * (Cell RE / Glom RE – 1).

### Immunoblotting

Immunoblots were performed from isolated glomerular cells from one mouse per lane. Samples were lysed in T-Per (Thermo Scientific) containing 1 mM sodium fluoride, 1 mM sodium vanadate, 1 mM calyculin A, complete (Roche) and denatured in 5x SDS. Samples were separated on a 4-12% Mini Protean TGX gel (BioRad, Hercules, CA) in a Tris-glycin migration buffer (0.25 M Tris, 1.92 M glycin, 1% SDS, pH 8.3). Protein transfer was performed in transfer buffer (0.192 M glycine, 25 mM Tris base, 20 % EtOH in ddH_2_O) in a TransBlot Turbo System (BioRad). After the transfer all proteins were visualized by Ponceau staining. PVDF membranes (Millipore) were blocked (5% nonfat milk) prior to incubation with primary antibodies diluted in Superblock blocking reagent (Thermo Scientific). Binding was detected by incubation with HRP coupled secondary antibodies (1:10000, 5% NFM). Protein expression was visualized with ECL SuperSignal (Thermo Scientific) according to manufacturer’s instructions on Amersham Imager 600 (GE Healthcare, Little Chalfont, UK). Western blots were analyzed using software from ImageJ (Abramoff M, 2004).

### Immunofluorescence

A 300 µm thick kidney slice from an mT/mG mouse was fixed with 4% PFA for 6 hours, washed with PBS and optically cleared for 24 hours using SCALEVIEW A2 (FUJIFILM Wako Chemicals). FACS-sorted glomerular cells were seeded on collagen Type IV (Sigma) coated 35 mm culture dishes (Sarstedt) for 24 hours, fixed with 4% PFA (EMSciences) for 8 min at RT. For immunofluorescence, unspecific binding was blocked with 5% normal horse serum in PBS (Vector) for 30 min at RT. Subsequently cells were stained with AF488-phalloidin (1:400, Molecular Probes), rhodamine-Wheat germ agglutinin (WGA, 1:400, Vector) and Hoechst (1:1000, Molecular Probes) for 30 min at RT and cover-slipped with Fluoromount (SouthernBiotech). Paraffin sections (3 µm) were deparaffinized and rehydrated to water. Antigen retrieval was performed by cooking at a constant 98°C in DAKO pH9 or citrate pH6.1 buffer for 30 min. Frozen sections were dried and fixed with 4% PFA for 8 min at RT. Unspecific binding in frozen and paraffin sections was blocked with 5% normal horse serum in 0.05% Triton X-100 (Sigma) for 30 min RT. Primary antibodies were incubated in blocking buffer o/n at 4°C: goat anti-THSD7A (1:200, Santa Cruz); Cy3-anti mouse IgG; rat anti-UCH-L1 (U104 1:50 (Sosna et al., 2013); guinea pig anti-nephrin (1:200, Acris); PE-podoplanin (1:50, BioLegend), AF647-CD73 (1:50, BioLegend); CD31 (1:100, PECAM1, BD Pharmingen). Following washes in PBS, fluorochrome labeled donkey secondary antibodies (all Jackson Immunoresearch Laboratories) and Hoechst (Molecular Probes) were applied where appropriate for 30 min at RT. After washes in PBS, sections were mounted in Fluoromount. Stainings were visualized with a LSM800 with Airyscan with the ZenBlue software (all Zeiss).

### UMAP Visualizations

To further visualize the separation of podocytes, mesangial and endothelial cells we performed Uniform Manifold Approximation and Projection (UMAP), a dimensionality reduction method (https://joss.theoj.org/papers/10.21105/joss.00861). This algorithm can be used to map high dimensional data into 2D. Visualizations were created from cytometry channel data and gates were used as labels for coloring. Only living single cells were included in analyses.

### Statistical analysis

Results were expressed as means ± SEM and significances were set at p < 0.05. The means were compared using the two-tailed nonparametric Mann-Whitney *U* test to enable robust conclusions on effects significance in case of departures from normality associated with small sample sizes. Replicates used were biological replicates, which were measured using different samples derived from distinct mice. More than two groups and timepoints were analyzed using the two-Way ANOVA with Sidaks multiple comparison.

## RESULTS

### Establishment of glomerular cell isolation method

First, we established and validated fluorescently labeled antibodies directed against extracellular antigens that would prove suitable for individual glomerular cell type labeling following an extensive protocol of enzymatic digestion and mechanical disruption (***Figure 1A***). Immunofluorescent analysis of frozen murine kidney sections (***Figure 1B***) demonstrated a specific labeling of podocytes with antibodies directed against podoplanin (Kabgani et al., 2012), of endothelial cells with antibodies directed against CD31 (Lertkiatmongkol et al., 2016), and of mesangial cells with antibodies directed against ecto-5′-nucleotidase/CD73 (e-5′NT/CD73), an enzyme responsible for adenosine formation from AMP expressed in mesangial cells (Castrop et al., 2004). Two protocols were used: As a reference for reporter-free glomerular live cell isolation, first mT/mG mice were used, in which podocytes are already marked by the intrinsic expression of eGFP and all the other murine cells by the intrinsic expression of tdTomato. To differentiate glomerular endothelial and mesangial cells, the remaining tdTomato positive glomerular cells were separated into tdTomato^+^/CD31^+^/eGFP^−^/CD73^−^ endothelial cells and into tdTomato^+^/CD73^+^/eGFP^−^/CD31^−^ mesangial cells (***Figure 1C***). This gating strategy was then modified for the use in wildtype mice using a combination of fluorophore-conjugated anti-podoplanin, anti-CD31, and anti-CD73 antibodies resulting in three distinct glomerular cell populations, namely podoplanin^+^/CD31^−^/CD73^−^ podocytes, CD73^+^/podoplanin^−^/CD31^−^ mesangial cells, CD31^+^/CD73^−^/podoplanin^−^ endothelial cells, and a population of non-labeled cells representing the contaminating tubular cells (***Figure 1D***). UMAP analysis demonstrates a clear separation of the three glomerular cell types in wildtype mice (***Supplemental figure 1A***).

### Isolated glomerular cells are pure

To evaluate purity and to assess further usability of the isolated cell populations, we performed several experiments. We first established the typical cellular yield obtained by this isolation technique (Fig. 2A). In total 11% (35,600 +/− 21000 SD) of glomerular cells isolated were podocytes, 16% (55,300 +/− 43,000 SD) mesangial cells, and 73% (245,000 +/ 136,000 SD) endothelial cells. These percentages and absolute cell numbers were stable across species and sex (***Supplemental figure 2A, 2B***). In old mice (61-69 weeks old), the efficiency of podocyte, mesangial and endothelial cell isolation decreased significantly, when compared to younger (11-14 weeks old) mice (***Supplemental figure 2C***). Percentages of isolated cell-types were not different between wildtype and mT/mG mice even though mT/mG mice provided a higher absolute number of podocytes, mesangial and endothelial cells than wildtype mice (***Supplemental figure 3***). We then plated the FACS-sorted cells onto collagen-type IV coated culture plates to assess the morphological characteristics of the different cell populations by visualization of the actin cytoskeleton with f-actin and of the glycocalyx by staining with wheat germ agglutinin (WGA) (***Figure 2B***). FACS-sorted podocytes were the largest cells and exhibited an elaborate phenotype with long arborizations and a cortical actin ring as expected (Pavenstadt et al., 2003). FACS-sorted mesangial cells were generally smaller than podocytes with a coarse actin cytoskeleton and smaller membrane arborizations. FACS-sorted endothelial cells were the smallest of all cell types, round in morphology with a prominent WGA positive glycocalyx. We then proceeded to analyze the relative expression of cell-specific transcripts in the FACS-sorted cell population relative to the transcript levels present in whole glomeruli from the same mouse (***Figure 2C***). The expression of the podocyte-specific transcripts *Nphs2* (encoding for podocin) and *Pdpn* (encoding for podoplanin) was enriched in podocytes compared to glomeruli and significantly lower in mesangial and endothelial cells. The expression of the mesangial cell-specific transcripts *Pdgfrb* (encoding for PDGFR-β) and *Cd73* (encoding for CD73) was enriched in mesangial cells compared to glomeruli and significantly lower in podocytes and endothelial cells. The expression of the endothelial cell-specific transcripts *Cdh5* (encoding for VE-Cadherin) and *Pecam1* (encoding for CD31) was enriched in endothelial cells compared to glomeruli and significantly lower in podocytes and mesangial cells. A potential contamination of our isolated cells with tubular or parietal epithelial cells was excluded by qPCR for the parietal cell specific transcripts *Lad1* (encodes for ladinin) and *Scin* (encodes for scinderin) (Kabgani et al., 2012); and for transcripts specific to tubular cells: *Aqp4* (encodes for aquaporin 4) and *Slc12a3* (encodes for sodium chloride channel NCC) (***Supplemental figure 4***). We next proceeded to evaluate the expression of cell-specific markers by immunoblot (***Figure 2D***). Cell-number-adapted lysates of FACS-sorted cells from individual mice were separated by SDS-PAGE and blotted for cell-specific markers. Corroborating the qPCR results, only podocytes expressed the slit membrane protein nephrin, only mesangial cells expressed the PDGFR-β, and only endothelial cells exhibited a VE-cadherin expression. Taken together, these analyses demonstrate the successful isolation of podocytes, glomerular endothelial and mesangial cells from individual mice in sufficient amounts to perform protein biochemical investigations.

**Figure 2:**
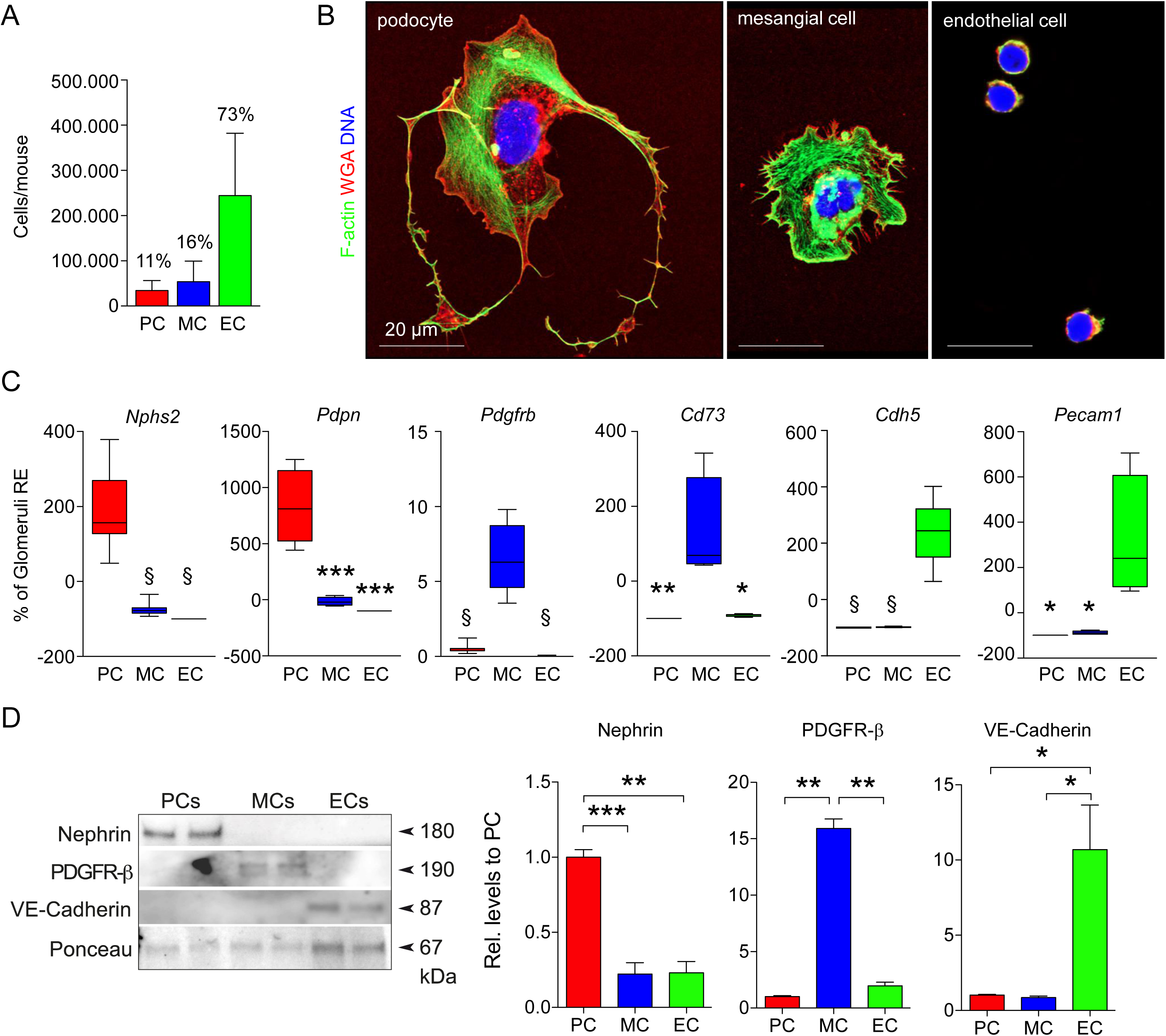
FACS-sorted cells are pure and in sufficient amounts to perform protein biochemical investigations. (**A**) Graph depicting the mean cell harvest per mouse in absolute numbers, mean +/−SEM, n=149, pooled data from 20 independent experiments. Percentages indicate the relative number of each cell population from the total amount of isolated cells. (**B**) Representative confocal images of FACS-sorted podocytes, mesangial cells and glomerular endothelial cells plated on collagen IV. F-actin (green) demarcates the actin cytoskeleton, Wheat germ agglutinin (red) the glycocalyx, DNA was stained with Hoechst (blue). (**C**) Real-time qPCR analysis exhibiting the expression of cell-specific transcripts. mRNA was isolated from FACS-sorted podocytes, mesangial and endothelial cells, and from isolated glomeruli derived from the same individual. ΔCT values were calculated using 18S as home keeper. ΔΔCT values were calculated as difference between glomerular and FACS-sorted cell type ΔCT. Displayed is the relative expression of FACS-sorted cell types in percent of the glomeruli. Note the significant enrichment of cell-specific transcripts in the FACS-sorted cell-types compared to the glomerulus, mean +/−SEM, \**p*<0.05; \*\**p*<0.01; \*\*\**p*<0.005; §*p*<0.0001, n=8 (*Nphs2, Pdgfrb, Cdh5*), n=4 (*Pdpn, Cd73, Pecam1*), pooled data from 3 independent experiments, one-way ANOVA and Bonferroni’s multiple comparisons test. (**D**) Cell-number-adapted lysates of FACS-sorted cells from individual mice were separated by SDS-PAGE and analyzed by immunoblot for the expression of cell-specific markers. Graphs exhibit densitometric quantification, values were normalized to ponceau of the same membrane and are expressed as mean +/−SEM relative expression levels to podocytes, \**p*<0.05; \*\**p*<0.01; \*\*\**p*<0.005, n>4 pooled data from 2-5 independent experiments.

### Proteomic analyses unravel glomerular cell specific molecular reference profiles and signaling networks

Using gating strategy 1 (applied to transgenic mice), we first generated a proteomics profiles from mT/mG mice using cell pellets consisting of one million cells. The dataset quantified 4753 proteins from these cells out of more than 6600 proteins. A 3-dimensional view of all these quantified proteins is given in (***Figure 3A***). Principal component analysis and hierarchical clustering revealed a strong separation of the cell-types (***Figure 3B***). Using t-test analyses, we defined lists of podocyte-specific, endothelia-specific and mesangial-specific proteins that were significantly enriched over the other cell-types. These can be found in ***Supplemental table 3*** for mT/mG mice and in ***Supplemental table 4*** for wildtype mice across sex, species, and age. We defined – based on statistical enrichment (at least two-fold log2 enrichment) followed by statistical testing with correction for multiple testing (p<0.05, FDR <0.01) 1150 endothelial specific, 366 mesangial specific, and 718 podocyte-specific proteins (FDR smaller than 0.01 and at least 2-fold log2 enrichment). To confirm the purity of the preparation, we mapped these cell type specific proteins against previously defined cell marker genes obtained by single-cell RNA sequencing (Karaiskos et al., 2018). The data was largely consistent with the protein expression defined on these clusters; however, some genes were differently expressed (***Figure 3C***). We mapped known proteinuria genes (Bierzynska et al., 2014) against the enrichment and found the majority of proteinuria genes in podocytes (***Figure 3D***). To further visualize the nature of these proteomes, we determined fold changes of enrichment factors of distinct uniprot key words. Radar plot analysis suggest different structural and metabolic functions of cell types, consistent with biology (***Figure 3E***). We used protein-protein interaction analyses and data resources to map potential inter-cellular protein crosstalk and communication. The analysis revealed contribution of different cell types to the extracellular matrix, as well as distinct signaling system. One example here was the ephrin-ephrin receptor system, with Epha2 and Efnb1 expressed in podocytes, and Epha2, Ephb4 and Efnb2 expressed in endothelial cells (***Figure 3F***). Furthermore, we constructed cell-type enriched signal transduction networks using a protein-protein interaction network and the NetBox algorithm (Cerami et al., 2010) (***Supplemental figure 5***). Both analyses support the view of podocytes as proteostatic active cells.

**Figure 3:**
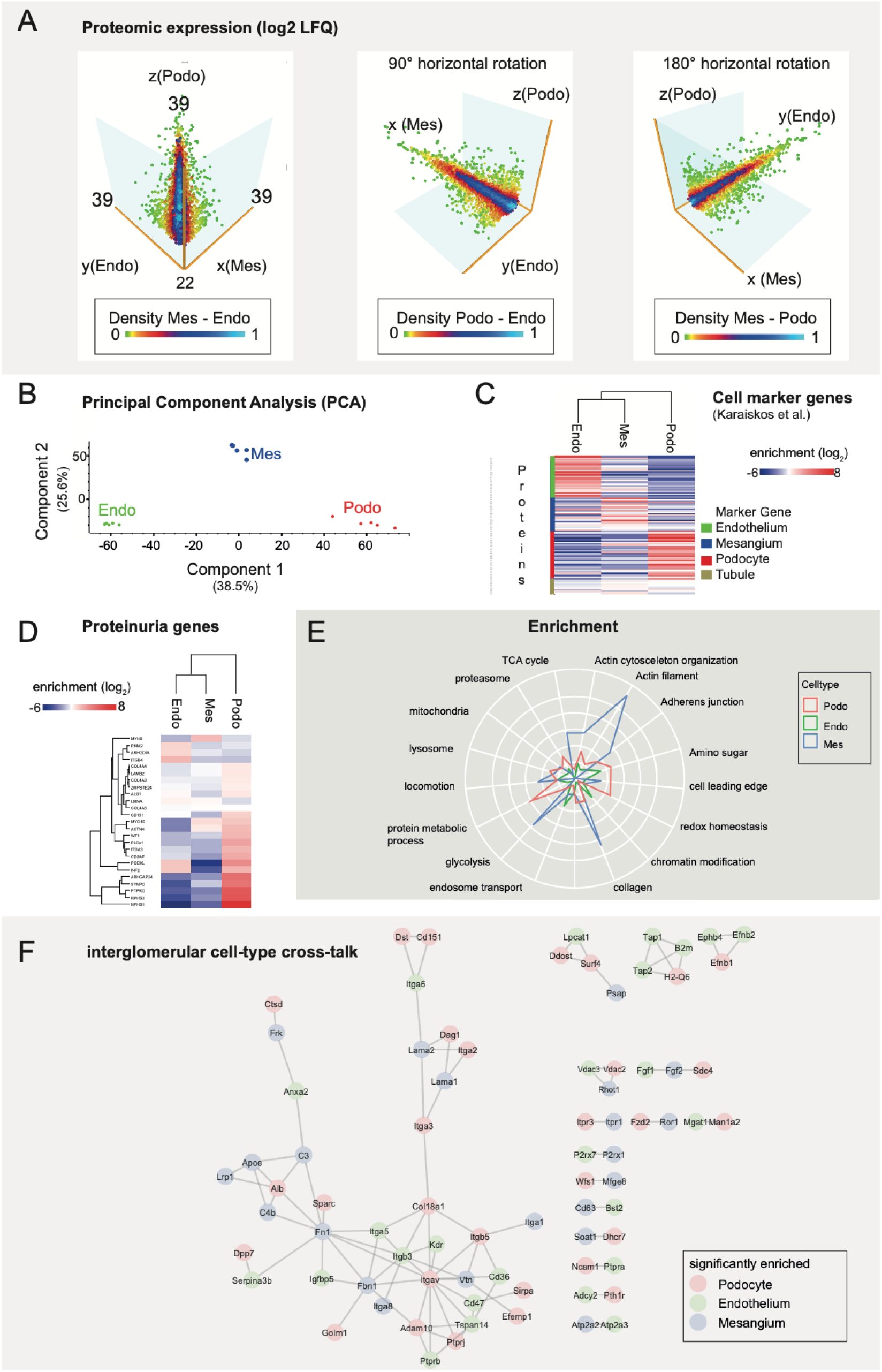
Tripartite cell-type isolation and proteomic analysis from mT/mG mice. (**A**) Threedimensional view of the tripartite cell proteomes containing 6600 proteins and quantification of 4400 proteins. (**B**) Principal component analysis (PCA) showing clear separations of endothelial, mesangial and podocyte proteomes. (**C**) Cell marker gene products derived from Karaiskos et al. (Karaiskos et al., 2018) and their proteomic expression from mesangial, podocyte and endothelial cells. (**D**) Expression of anotated proteinuria genes. (**E**) Radarplot showing log2 enrichments of different GO terms in mesangial, podocyte and endothelial cells. The center shows a fold change of 0, and every circle is a step of 2 fold enrichment in this particular population. (**F**) Visualization glomerular cell cross-talk systems. Protein-protein interaction networks were generated in STRING using high-confidence interactions (database and experimental evidence, <0.9).

### Proteomic analyses unravel new glomerular cell specific proteins

We applied the newly generated methods to cell populations from a single mouse, comparing male and female mice in a C57BL/6 background, BALB/c females and – as control – mT/mG female mice that were subjected to the original protocol. To test the generalizability of the method from FACS-sorted cells, and to test the purity of the preparations, we generated proteomics profiles from three different cell types by single-shot proteomics from the cell fractions from a single mouse. An overview of the dataset revealed very clear separations of three distinct cell types, suggesting that the identity of these cells is in fact different (***Figure 4A***). We again calculated then glomerular specific enriched proteins, finding novel markers of the cell types by comparison with the human protein atlas (***Figure 4B, Supplemental figures 6, 7, 8***). To this end, and to the best of our knowledge we identified 12 new podocyte-enriched proteins, 16 new mesangial cell-enriched proteins, and 22 new glomerular endothelial cell-specific proteins based on strict criteria. In addition, the clear presence of mesangial cells in this protocol is remarkable, since the molecular protein composition of these cells has not been described. To our astonishment our proteomic data, qPCR, and immunofluorescent validation experiments restricted β3-integrin and its ligand fibronectin expression to glomerular endothelial cells (***Supplemental figure 9***), and not to podocytes in mice, contradicting published reports for the expression of both proteins in murine podocytes *in vivo* (Kliewe et al., 2019; Wei et al., 2011; Wei et al., 2008). We next analyzed, which protein classes had high variance between the four strains (***Figure 2C***). Podocyte proteins had the most variance (highest standard deviation) between the proteins among all the three cell types, a fact that may be due to the disruptive isolation method used in this protocol. Through the high sensitivity of the assay, we also observed that the mesangial cell populations had a minimal contamination with podocytes with the initial gating strategy (***Supplemental figure 1B, 1C, and 10A***) that we estimated to be around 2% as compared to the mT/mG-derived mesangial cells. In reaction to this observation, we improved the gating strategy for all other investigations (***Figure 1D***). UMAP analysis demonstrates the removal of the contaminating podocytes in the mesangial cell population with the adjustments (***Supplemental figure 1B, C***).

**Figure 4:**
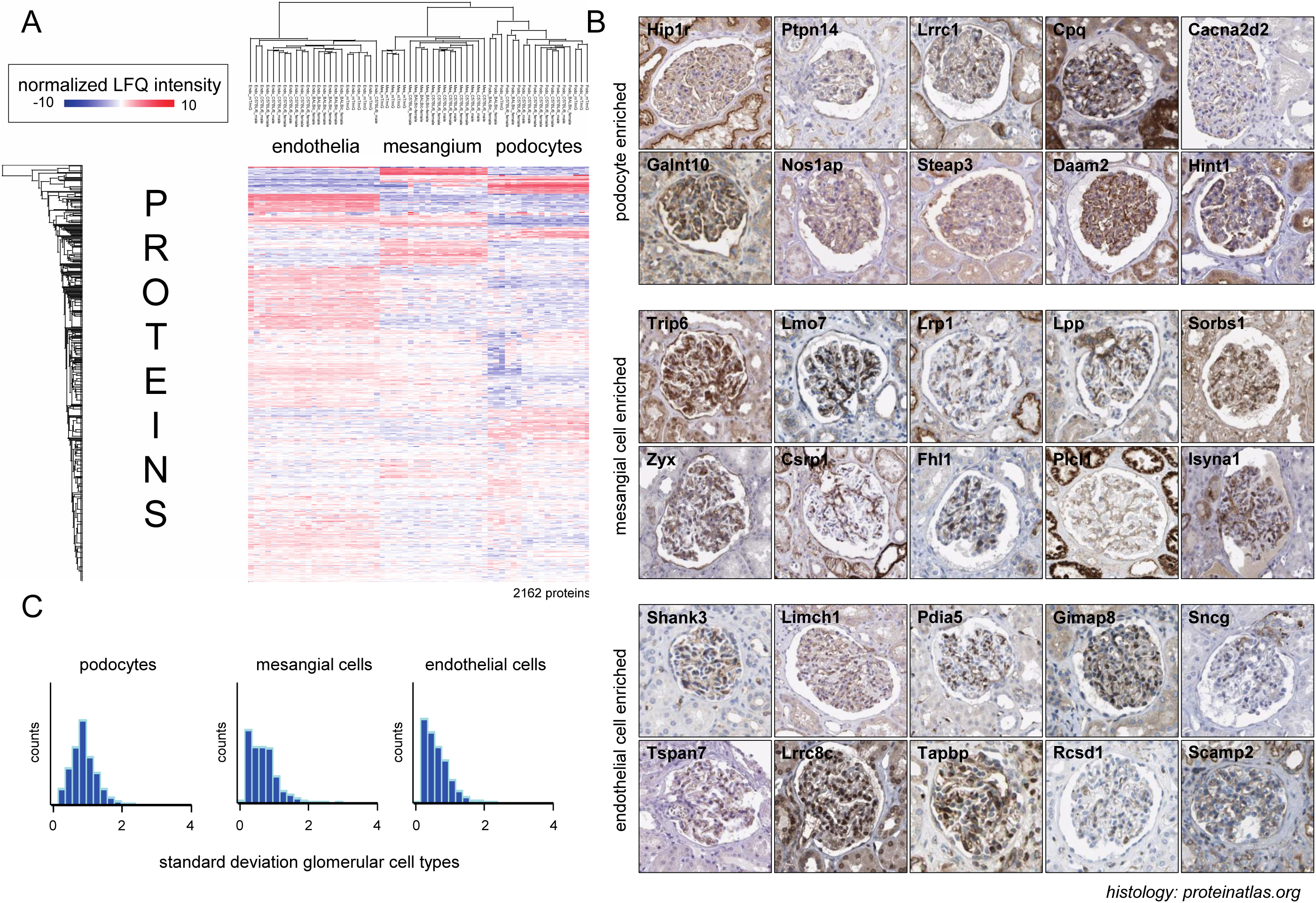
Identification of new podocyte, mesangial and glomerular endothelial cell enriched proteins from tripartite cell-type proteomics from reporter-free single mice. (**A**) Heatmap showing euclidean distance clustering of single-mouse single shot proteomics results from male C57BL/6, female C57BL/6, female BALB/c and female mT/mG mice. Each columns are cell-type proteomes from a single mouse. (**B**) To identify new proteins enriched in podocytes, mesangial cells and glomerular endothelial cells, glomerular cell-types proteome lists of individual mice were compared. For the comparisons we defined 1) a Student t-test difference cut-off of > 2 between podocytes and non-podocytes, 2) a negative search result in pubmed (https://www.ncbi.nlm.nih.gov/pubmed) and and 3) a validated podocyte expression pattern in the human protein atlas (https://www.proteinatlas.org/search), from which the histological micrographs are taken from. (**C**) Standard deviation distributions of proteins in each cell-types reveals low variance of expression in endothelial cells.

### Glomerular endothelial cells exhibit proteomic differences between mouse strains

We performed deeper proteomic bioinformatic analysis of the population with the most homogeneity, the glomerular endothelial cells. Principal component analysis of the glomerular endothelial cell proteomes derived from four different strains female C57BL/6, male C57BL/6, BALB/c, and mT/mG mice was performed. Data revealed a separation by strain, with male and female C57BL/6 being similar, and mT/mG being most different from these (***Figure 5A***). UCH-L1 (Ubiquitin C-terminal hydrolase L1, a major neuronal deubiquitinates of the degradative ubiquitin proteasome system (Bishop et al., 2014)) and Gvin1 (GTPase, very large interferon inducible 1, which contributes to the cellular response to both type I and type II interferons (Klamp et al., 2003)) were proteins that contributed the most to this dataset (***Figure 5B***), suggesting that reporter-dependent artefacts in the proteome must be considered in the use of fluorescent reporter systems. Volcano plot analysis revealed a more than 16-fold increase of these proteins in the mT/mG endothelial cells as compared to C57BL/6 cells (***Figure 5C***). Exemplarily, the differences in endothelial expression of UCH-L1 could be confirmed using confocal microscopy of C57BL/6 and mT/mG glomeruli (***Figure 5D***). We also performed unbiased analysis of differentially expressed proteins within the four strains (***Figure 5E***), ANOVA with FDR <0.01. Clustering analysis revealed that there was a significant alteration in stress-response proteins in mT/mG mice compared to the other strains and an alteration of immunity and cell adhesion proteins in BALB/c females compared to C57BL/6 females (***Figure 5F***).

**Figure 5:**
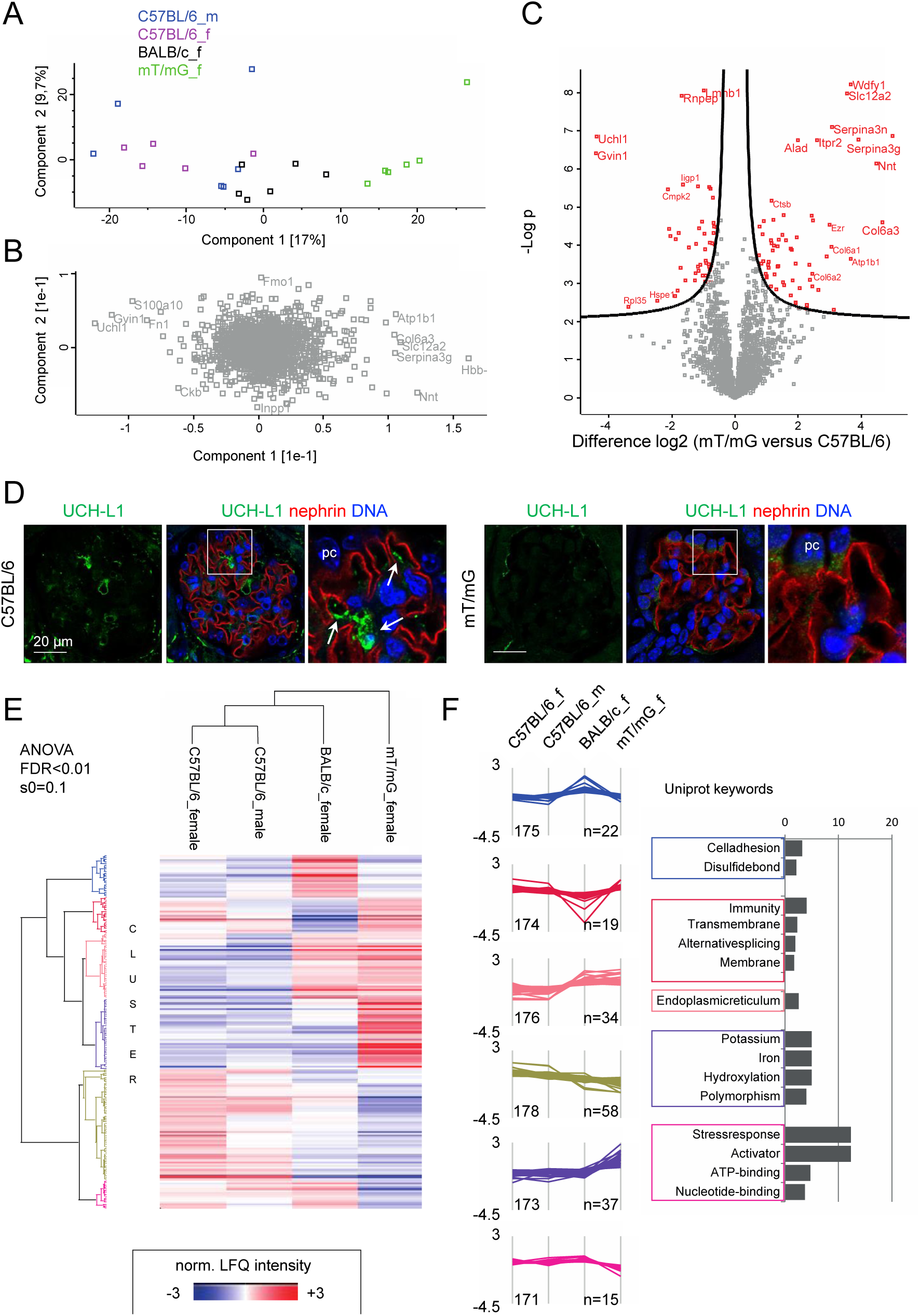
Strain-dependent differences in endothelial proteomes. (**A**) Analysis of glomerular endothelial proteome from wildtype mouse. (**B**) Principal component analysis of proteins. (**C**) Volcano plot showing differences between proteins highly expressed in mT/mG (right side) and C57BL/6 mice (left side) of the plot. (**D**) Representative confocal images of UCH-L1 (green) expression in a C57BL/6 and in a mT/mG glomerulus. Nephrin (red was used to demarcate the glomerular filtration barrier, DNA (blue). Arrows point toward endothelial UCH-L1 expression, pc = podocyte. (**E**) Hierarchical clustering of mean normalized protein expression (log2 LFQ) from four different mouse strains. (**F**) Clustering combined with GO term enrichment reveals stress response proteins in endothelia from mT/mG mice.

### Glomerular cell types resolve specific reactions in anti-THSD7A membranous nephropathy

We proceeded to analyze, whether our glomerular cell isolation technique was also suitable for the separation of glomerular cell types in the setting of glomerular damage/stress/inflammation. For this purpose, we induced defined glomerular injury through injection of rabbit antibodies, which specifically target the podocyte foot-process antigen THSD7A (Herwig et al., 2019). This immunological model of THSD7A-associated membranous nephropathy (Tomas et al., 2017) leads to selective podocyte injury with massive proteinuria within days, and mimics the pathophysiology of human THSD7A-associated membranous nephropathy (Tomas et al., 2014). CD45 as an additional marker was added to the FACS-sort protocol to remove potential leukocytes adhering to the endothelium of the perfused glomeruli in inflammation (***Supplemental figure 11***). A UMAP demonstrated separation of CD45-positive cells from all other cells (***Supplemental figure 1D***). No significant infiltration and adherence of leukocytes could be detected in this glomerulonephritis model (***Supplemental figure 12A, B***). Further, the cell sort marker expression of podoplanin, CD31, and CD73 was mostly preserved in day 7 glomeruli of highly proteinuric mice (***Supplemental figure 12C***), a predisposition for the successful isolation of podocytes, glomerular endothelial, and mesangial cells. Glomerular cell-types were analyzed for a stress response by qPCR on day 1 after antibody binding prior to development of proteinuria and on day 7 when severe podocyte injury and nephrotic syndrome were established (***Figure 6A and B***). To this end, qPCR analyses demonstrated significant changes in podocyte gene transcription as early as day 1 for 1) genes involved in glomerular basement membrane (GBM) remodeling (***Figure 6C***: agrin (*Agrin)*, a major proteoglycan of the GBM (Groffen et al., 1998); laminin chain β1 (*Lamb1*), laminin chain of the immature / injury remodeled GBM (Abrahamson et al., 2003)); 2) for genes involved in an oxidative stress response (***Figure 6D***: vascular endothelial growth factor a (*Vegfa*), involved in angiogenesis and maintenance of the glomerular endothelium (Eremina et al., 2007); superoxide dismutase 1 (*Sod1*), detoxifies reactive oxygen radicals (Sea et al., 2015)); and 3) for genes of key metabolic enzymes (***Figure 6E***: pyruvate kinase (*Pkm*), key enzyme of glycolysis (Gupta and Bamezai, 2010); acyl-CoA dehydrogenase, long chain (*Acadl*), key enzyme in β-oxidation of fatty acids (Chegary et al., 2009) in mice exposed to rbTHSD7A-abs in comparison to control rbIgG. Importantly, these changes in gene transcription were not visible upon analyses of isolated glomeruli on day 1 and day 7, demonstrating the enhanced sensitivity obtainable by performing glomerular cell-type-based analyses. Mesangial and glomerular endothelial cells showed no major alterations in evaluated transcription levels on day 1 or day 7. Strikingly, the observed gene induction on day 1 was not apparent on day 7 anymore in rbTHSD7A-abs injected mice. On day 7 all analyzed transcript levels were repressed in rbTSHD7A-MN podocytes in comparison to day 1 and only slightly elevated in comparison to day 7 rbIgG injected control mice (data not shown), suggesting a specific and acute reaction to rbTHSD7A antibodies on day 1.

**Figure 6:**
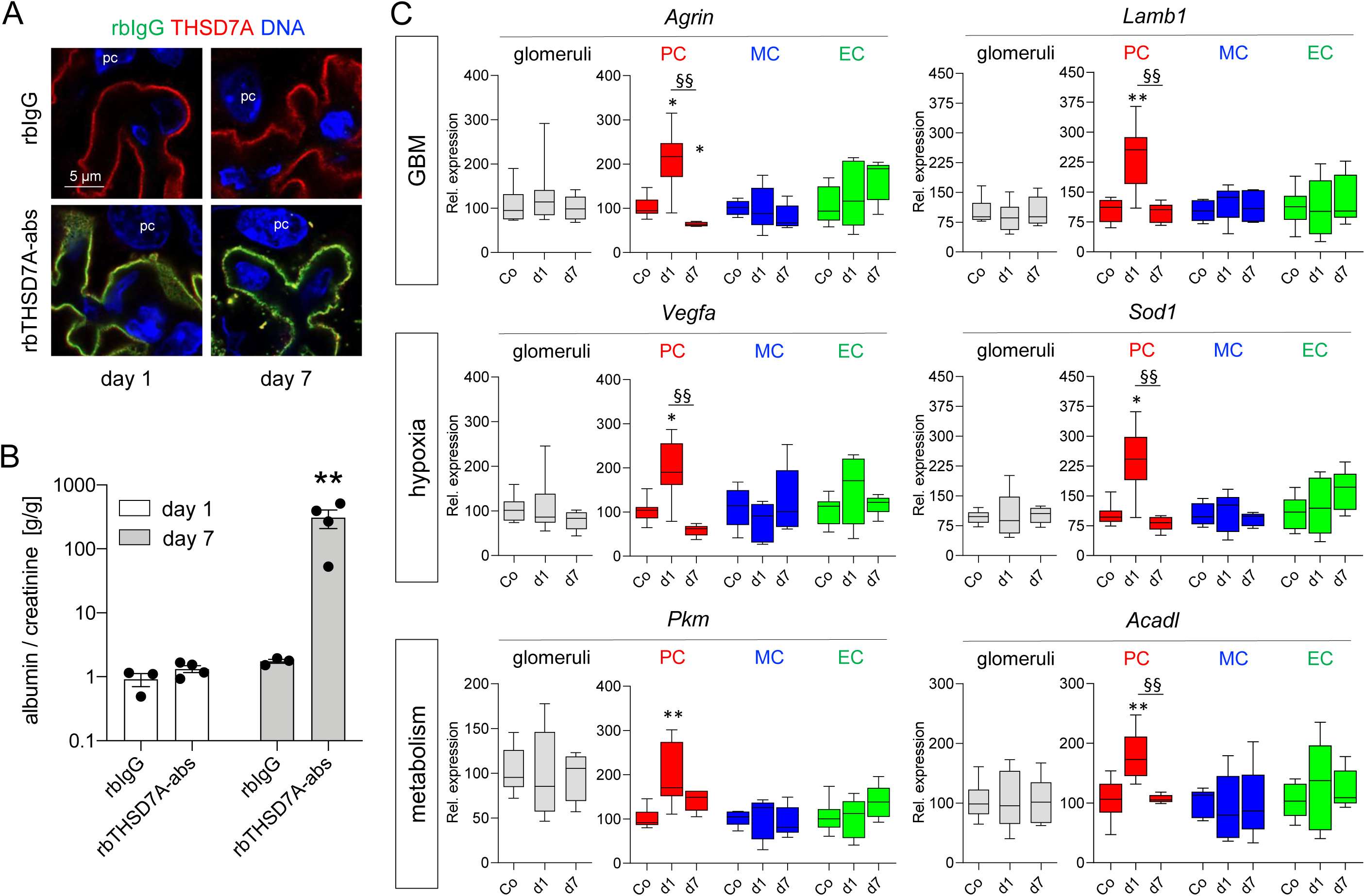
Glomerular cell specific analyses in the experimental model of THSD7A membranous nephropathy unravels acute transcript regulation. Experimental THSD7A membranous nephropathy was induced in male BALB/c mice by injection of podocyte-targeting rabbit anti-THSD7A antibodies (rbTHSD7A-abs) or unspecific control rabbit IgG (rbIgG). Analyses were performed on day 1 and 7 after antibody injection. (**A**) Confocal analyses demonstrating rabbit IgG (rbIgG, green) deposition in a linear (day1) and a more granular (day 7) pattern in rbTHSD7A-abs injected mice. Yellow colour exhibits colocalization of the injected rabbit antibodies with the antigen THSD7A (red), as a sign for successful induction of membranous nephropathy, DNA (blue), pc = podocyte. (**B**) Urinary albumin to creatine ratio on days 1 and 7. \*\**p*<0.01 to day 7 rbIgG, Mann Whitney *U* test. (**C**) qPCR analyses were performed in isolated glomeruli (as the conventional method, graphs with grey bars) and in FACS-sorted (right graphs) podocytes (PC, red bars), glomerular endothelial (EC, green bars) and mesangial cells (MC, blue bars) of the same mouse. Graphs exhibit relative expression in comparison to rbIgG treated mice (control, Co, stripped bar) of the respective time point (day 1 rbIgG to day 1 rbTHSD7A-abs and day 7 rbIgG to day 7 rbTHSD7A-abs) after normalization to 18S as homekeeper. The control (Co) bars exhibit the pooled day 1 and day 7 rbIgG values for simplification. Mann Whitney *U* test, \**p*<0.05, \*\**p*<0.01 to control, §§*p*<0.01 to day 7 rbIgG.

## DISCUSSION

The complicated architecture of the kidney hampers the direct access of susceptible cell types in kidney diseases for further functional analysis. Here, we established a generalizable method for generating molecular reference profiles of all kidney glomerular cell types in health and disease. This FACS-sort based method, which we propose to name timMEP (tripartite <u>i</u>solation method of <u>m</u>urine <u>M</u>esangial, glomerular <u>E</u>ndothelial cells, and <u>P</u>odocytes) results in the precise isolation of the three glomerular cell types as extensively validated by qPCR, proteomic, immunoblot and immunofluorescent analyses. TimMEP has the immense advantage of not requiring the extensive breeding of transgenic reporter mouse models for subsequent glomerular cell type isolation, of saving time / animals / money and of preventing reporter-induced artifacts. Hence, timMEP is theoretically amenable to any existing mouse model and will facilitate and sharpen investigations to glomerular cell biology in a cell-type specific, cost-effective and timely manner.

Despite these huge advantages, timMEP has limitations. A key requirement for timMEP is a maintained expression and accessibility of the cell-type specific extracellular protein epitopes to fluorophore-conjugated antibodies for FACS-sorting. Frequently, glomerular cells change their proteome and thereby cell type marker expression in the setting of stress (Koehler et al., 2020; Meyer-Schwesinger et al., 2009), a process often reflecting a dedifferentiation (especially of podocytes (Guhr et al., 2013)) or inflammatory activation (especially within endothelial and mesangial cells) of glomerular cells. In addition, proteolysis occurs extracellularly to large extents (Rinschen et al., 2018b). We therefore carefully selected glomerular cell-type markers, which were in our hands relatively stable expressed by the glomerular cell types in health and disease and provide other cell-type markers that could be used for modified FACS-sorting (***Supplemental table 5***). In line, we were able to isolate podocytes, mesangial and glomerular endothelial cells in the murine antigen specific model of THSD7A membranous nephropathy. This succeeded both shortly after disease induction (day 1) and in established massive proteinuria (day 7). In these settings, the proof-of-principle investigations demonstrated the advantage of timMEP: Early podocyte specific effects of the injected anti-THSD7A antibodies were discernible, which would have been missed by conventional analyses of isolated glomeruli.

A second limitation of timMEP is the long isolation procedure, which comprises 1) glomerular isolation with magnetic beads, 2) glomerular single cell preparation by digestion with a protease cocktail, and by mechanical disruption at 37°C, and 3) mechanical stress in the FACS-sorter. It therefore needs to be taken into consideration that, together, these inevitable factors might ultimately change the transcriptome and proteome of glomerular cells. This experimental artifact however needs to be considered for any kind of cell isolation/sample generating procedure and could be mitigated by adequate controls and experimental design. Despite these drawbacks, the amount of cell debris and of dead cells detected by FACS after timMEP was low. We were able to culture viable podocytes, mesangial cells and glomerular endothelial cells suggesting that the strain practiced on the glomerular cell types was present but not detrimental.

The number of isolated podocytes, mesangial and glomerular endothelial cells per mouse using timMEP is sufficient for routine qPCR analysis, transcriptomic and proteomic analyses. Immunoblot analyses of isolated podocytes, mesangial and glomerular endothelial cells of individual mice are also feasible against highly abundant proteins, as shown by the immunoblot against the cell type specific proteins nephrin, VE-cadherin, and PDGFR-β. However, protein analyses of less abundant proteins will in some cases require pooling of glomerular cell types isolated by timMEP from 2-4 mice to reach a sufficient protein concentration for the analyses. The efficiency of timMEP is best in mice aged 6-20 weeks, thereafter the number of isolated podocytes, mesangial and glomerular endothelial cells decreases. This might partly be based on the fact that the efficiency of glomerular isolation decreases with age and that the number of glomerular cells (especially of podocytes) decreases with age (Wanner et al., 2014), or aged cells might be more vulnerable. However, we could not state a rise in cell debris or dead cells following timMEP in aged mice.

Using timMEP, the presented dataset provides for the first time a reference for the proteomes of the three cell types in the glomerulus. While we already described the podocyte proteome (Rinschen et al., 2018a), this dataset now also covers cell types that are not easily amenable to single-cell dissociation, such as mesangial cells at a reasonable depth of 6300 proteins. Single cell transcriptomics data generally have a large under-representation of mesangial cell capture as compared to what is expected based on morphometry (Clark et al., 2019). Analysis of several single cell-datasets revealed that mesangial cells are not “easily accessible” given their common renin lineage (and thus the chance of misclassification). Our dataset, in contrast, now provides unambiguous mRNA and protein-based markers that can be used to also benchmark single-cell datasets regarding the presence of this particular cell type and that is consistent with the few mesangial cells (around 2%) in a previous single RNAseq cell dataset derived from glomerular populations (Karaiskos et al., 2018). As a result, we indeed could identify 13 new enriched podocyte proteins, 16 new enriched mesangial cell proteins, and 23 new enriched glomerular endothelial cell proteins using this method in naïve wildtype cells. Further, our analyses unraveled a strain and transgene-dependent expression of proteins among glomerular cell types offering insights into protein machineries among strains. Thereby, the proteome of endothelial cells was significantly different between BALB/c, C57BL/6 and mT/mG mice, possibly one explanation to the differential susceptibility of different mouse strains to glomerular disease models (Meyer et al., 2007; Meyer-Schwesinger et al., 2011; Tomas et al., 2017). These observations further underline the differences of mouse strains commonly used in renal research, already stated by Randles et al. for the strain dependent composition of the glomerular basement membrane (Randles et al., 2015). Moreover, these findings strongly suggest the necessity to consider the presence of reporter-associated artefacts in the use/ interpretation of transgenic reporter mice, as the expression of fluorescent reporters such as eGFP constitutes up to 6% of the proteome (Rinschen et al., 2018a), a fact that could affect distinct cellular pathways such as degradation pathways. In line, UCH-L1, a major deubiquitinating enzyme of the neuronal proteasomal degradation system was one of the most differentially regulated proteins between the endothelia of mT/mG and C57BL/6 mice. Finally, the data can be mined for protein-ligand receptor interactions and factors of intracellular communications. The ephrin-ephrin receptor system was highlighted here, suggesting podocyte-podocyte or/and podocyte-endothelial cross-talk. Epha2 has not been localized to glomerular cells yet (our analyses detected Epha2 on podocytes and glomerular endothelial cells), however, an involvement in the physiology of the renal tubular system was proposed (Xu et al., 2005). We found the expression of the Ephb4 – Efnb2 system in adult endothelial cells, an expression described in the context of the developing kidney (Takahashi et al., 2001) and in the recovery from glomerulonephritis (Ephb4 on podocytes (Wnuk et al., 2012)). In line with our findings, localization of Efnb1 on podocytes has been reported (Hashimoto et al., 2007), and an involvement in diabetic nephropathy through an interaction with Partitioning-defective (Par)-6 (Takamura et al., 2020) and in JNK pathway regulation (Fukusumi et al., 2018) have been demonstrated.

In summary, we here present a generalizable and time/cost/animal effective method to isolate glomerular cell types, which will enable a widespread and detailed glomerular cell-type resolved transcriptional and protein biochemical analysis to deepen our future understanding of glomerular cell biology in health and disease.

## ACKNOWLEDGEMENTS

We are grateful to the staff of the FACS Sorting Core Unit for excellent technical assistance. FH and UW developed the glomerular cell isolation technique, performed the experiments and analyzed the data. SS performed the primary cell culture experiments. MMR performed the proteomic analyses. TBH provided the mT/mG mice. WS and MS performed immunoblot analyses. CMS developed the glomerular cell isolation technique, performed immunofluorescence and confocal microscopy, analyzed the data. FH, UW, MMR and CMS wrote the manuscript. This work was supported by the DFG (SFB1192, project B3 to CMS, B8 to TBH, DFG Ri2811/1 and DFG Ri2811/2 to MMR) and Else-Kröner Fresenius Stiftung (“Else Kröner-Promotionskolleg Hamburg – Translationale Entzündungsforschung” (iPRIME). Work in M.M.R.’s lab is supported by the Young Investigator Award by the Novo Nordisk Foundation, grant number NNF19OC0056043.

