## Supplemental appendix for "Tripartite separation of glomerular cell-types and proteomes from reporter-free mice"

***Supplemental table 1:*** Antibodies and dyes used in the study for FACS sort.

***Supplemental table 2:*** Primers used in the study.

***Supplemental table 3:*** Podocyte-specific, endothelia specific and mesangial specific proteins that were significantly enriched over the other cell types in mT/mG mice.

***Supplemental table 4:*** Podocyte-specific, endothelia specific and mesangial specific proteins that were significantly enriched age, sex, and species in wildtype mice.

***Supplemental table 5:*** Alternative membrane proteins potentially useful for timMEP in diseased glomeruli.

***Supplemental figure 1:*** UMAP analysis of FACS sorted glomerular cells.

***Supplemental figure 2:*** Counts of isolated cells were comparable in between gender, species but not age.

***Supplemental figure 3:*** Comparison of FACS-sort between mT/mG mice and wildtype mice.

***Supplemental figure 4:*** FACS-sorted glomerular cells are not contaminated with parietal epithelial or tubular cells.

***Supplemental figure 5:*** Tripartite cell type isolation and proteomic analysis from mT/mG mice.

***Supplemental figure 6:*** Identification of new podocyte enriched proteins.

***Supplemental figure 7:*** Identification of new mesangial cell enriched proteins.

***Supplemental figure 8:*** Identification of new glomerular endothelial cell enriched proteins.

***Supplemental Figure 9:*** Integrin-$\beta$3 and its ligand fibronectin 1 are predominantly expressed on glomerular endothelial cells.

***Supplemental figure 10:*** Comparison of old and new gating strategy.

***Supplemental figure 11:*** Gating strategy used for the separation of glomerular cell types in the mouse model of anti-THSD7A membranous nephropathy.

***Supplemental figure 12:*** Leukocytes are not increased in glomeruli in the setting of experimental THSD7A-associated membranous nephropathy.

| Antibody | Clone | Company | Dilution |
|---|---|---|---|
| PE-Podoplanin | 8.1.1 | BioLegend | 1:200 |
| AF647-CD73 | TY/11.8 | BioLegend | 1:2000 |
| BV421-CD31 | MEC 13.3 | BD OptiBuild | 1:800 |
| AF700-CD45 | 30-F11 | BioLegend | 1:100 |

| Dye | Company | Dilution |
|---|---|---|
| Near-IR fluorescent reactive dye | Invitrogen | 1:4000 |

***Supplemental table 1:*** Antibodies and dyes used in the study for FACS-sort.

| Target | Sequence 5‘ – 3‘ |
|---|---|
| *Agrn* -forward | CTAGGGGAATCTCCGGTCCC |
| *Agrn* -reverse | GGCCTCTTAGAGACACCAGC |
| *Aqp4* -forward | CCCGCAGTTATCATGGGAAAC |
| *Aqp4* -reverse | GAAGGCTTCCTTAAGGCGACG |
| *Cdh5* -forward | CGTTGGACTTGATCTTTCCC |
| *Cdh5* -reverse | CGCCAAAAGAGAGACTGGAT |
| *Fn1* -forward | CTTGCACGATGATATGGAGA |
| *Fn1* -reverse | AGCTGAACACTGGGTGCTAT |
| *Hk1* -forward | CGGAATGGGGAGCCTTTGG |
| *Hk1* -reverse | GCCTTCCTTATCCGTTTCAATGG |
| *Itgb3* -forward | GCGCGCGGCGTGAAG |
| *Itgb3* -reverse | CACGCCTCGTGTGGTACAGAT |
| *Lad1* -forward | CACAGCCATCCAGAGGTCAG |
| *Lad1* -reverse | TCAAAGAGGTGTCGCTTGCT |
| *Lamb1* -forward | TCCCCGCTACCTCTCCAGAA |
| *Lamb1* -reverse | CCATAGGGCTAGGACACCAAA |
| *Nphs2* -forward | TGACGTTCCCTTTTTCCATC |
| *Nphs2* -reverse | AGGAAGCAGATGTCCCAGT |
| *Nt5e* -forward | GCAGCATTCCTGAAGATGCG |
| *Nt5e* -reverse | CTCCCGAGTTCCTGGGTAGA |
| *Pdgrfb* -forward | TGGTATCACTCCTGGAAGCC |
| *Pdgrfb* -reverse | AACAGAAGACAGCGAGGTGG |
| *Pdpn* -forward | GGGATGAAACGCAGACAACAG |
| *Pdpn* -reverse | TTTAGGGCGAGAACCTTCCAG |
| *Pecam1* -forward | AGCCAACAGCCATTACGGTTA |
| *Pecam1* -reverse | AGCCTTCCGTTCTCTTGGTG |
| *Pkm* -forward | GCCGCCTGGACATTGACTC |
| *Pkm* -reverse | CCATGAGAGAAATTCAGCCGAG |
| *Scin* -forward | CCAGAATTGTGGAGGTTGACG |
| *Scin* -reverse | GCCATTGTTTCGTGGCAGTT |
| *Slc12a3* -forward | CGGGATAATGGCAAGGTCAAGTCG |
| *Slc12a3* -reverse | GGAATTCTGATGCGGATGTCATTGATGG |
| *Sod1* -forward | GGAACCATCCACTTCGAGCA |
| *Sod1* -reverse | CTGCACTGGTACAGCCTTGT |
| *Vegfa* -forward | AATGCTTTCTCCGCTCTGAA |
| *Vegfa* -reverse | GCTTCCTACAGCACAGCAGA |

***Supplemental table 2:*** Primers used in the study.

| Enriched in podocytes | Enriched in mesangial cells | Enriched in glomerular endothelial cells |
|---|---|---|
| Kirrel1 | Itga8 | Dysf |
| Enpep | Gja5 | Emcn |
| Ptpro | Lrp1 | Tmem2 |
| Cxadr | Mcam | Gimap1 |
| Galnt10 | Cspg4 | Slc43a3 |
| Sdc4 | Col6a2 | Kdr |
| Crb2 | Col6a1 | Tspan7 |
| Itgb5 | | Ece1 |
| Itm2b | | Ppap2a |
| Cldn5 | | Lpcat1 |

***Supplemental table 5:* Alternative membrane proteins potentially useful for timMEP in diseased glomeruli.** The used FACS-sort markers presented here for timMEP (podoplanin, CD73 and CD31) might be altered in specific glomerular disease models. Therefore, we analyzed our proteomic data for possible alternatives. The presented proteins are enriched in the respective glomerular cell type and have been annotated to reside on the cellular membrane. Cell type-specific expression has been verified in the Human Protein Atlas. If an alternative to our standard targets is needed, this list may serve as a starting point.

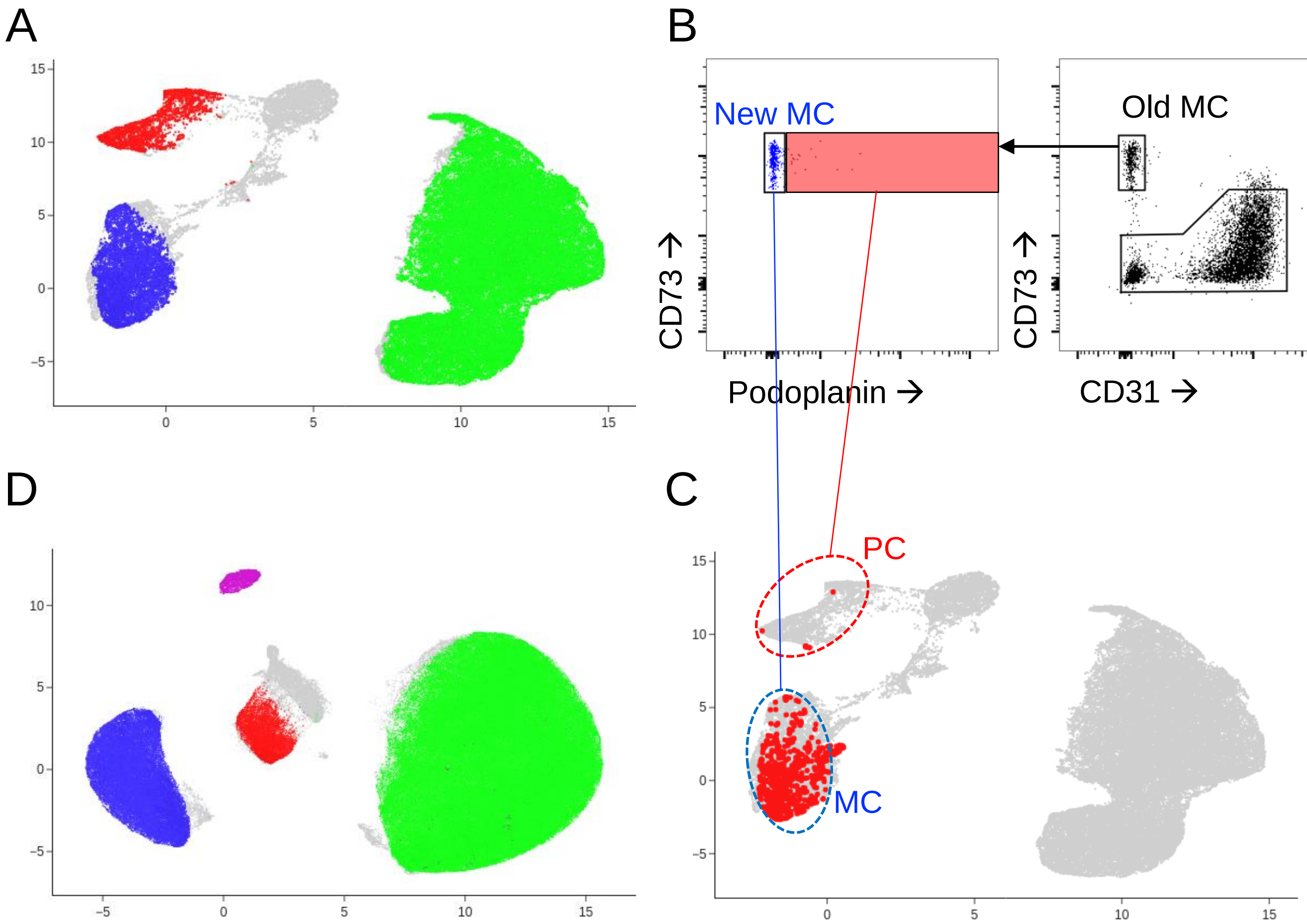


***Supplemental figure 1:*** **UMAP analysis of FACS-sorted glomerular cells. (A)** UMAP plot of cytometry data recorded during sorting of wildtype cells used for proteomic analyses in this paper. The plot demonstrates clear clustering of podocytes (PC, red), mesangial (MC, blue) and endothelial cells (green), respectively. **(B)** Modified snippet taken from Fig. 1D demonstrating the differences between the earlier and revised gating strategy. Highlighted in the light red box are cells that might have caused a slight podocyte contamination in the mesangial cell population. **(C)** To further investigate actual contamination we overlaid these cells (light red box in B) in red on to the UMAP shown in A. Interestingly, most of those cells reside inside the mesangial cell cluster (blue dotted ellipse; previously correctly classified as mesangials) with only very few cells residing in the podocyte cell cluster (red dotted ellipse in A; previously misclassified as mesangials). Furthermore, the improved gating strategy removed the contamination as there are no mesangials (blue dots) in the podocyte cluster (red dots) in A. **(D)** UMAP of cytometry data with additional CD45 gating to visualize immune cells (violet cluster).

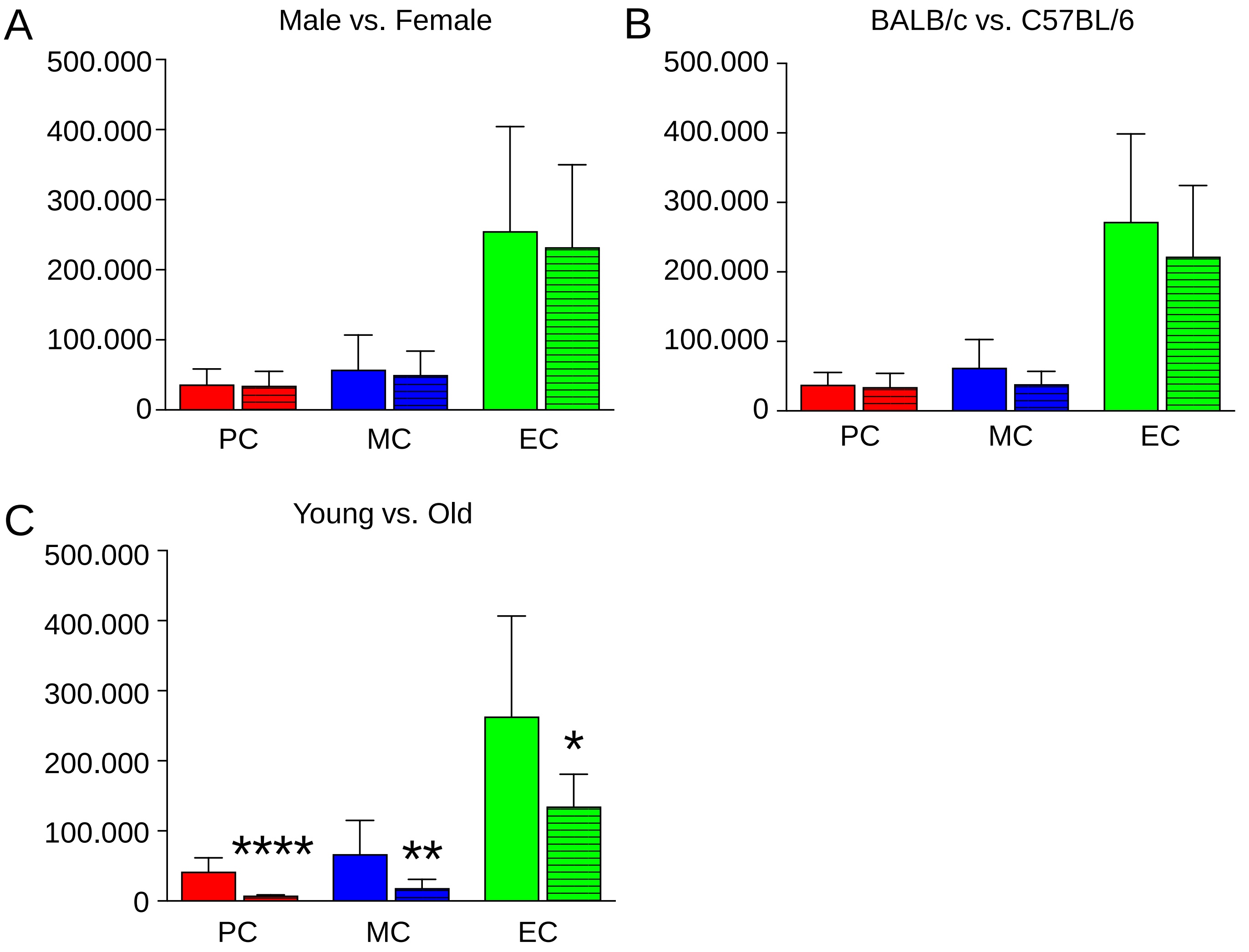


***Supplemental figure 2:*** **Counts of isolated cells were comparable in between gender, species but not age.** Absolute number of podocytes (PC), mesangial (MC), and endothelial cells (EC) FACS-sorted from the isolated glomerular cells per mouse, mean +/-SEM, n=5-74, pooled data from 18 independent experiments. **(A)** Empty bars represent cell counts from male C75BL/6 mice, stripped bars from female C75BL/6 mice, all aged 11-14 weeks. **(B)** Empty bars represent cell counts from female Balb/C mice, stripped bars from female C75BL/6 mice, all aged 11-14 weeks. **(C)** Empty bars represent cell counts from young male C75BL/6 mice 11-14 weeks of age, stripped bars from old male C75BL/6 mice aged 61-69 weeks. *$p<0.05$, **$p<0.01$, ****$p<0.0001$ to young male mice, Mann Whitney U test.

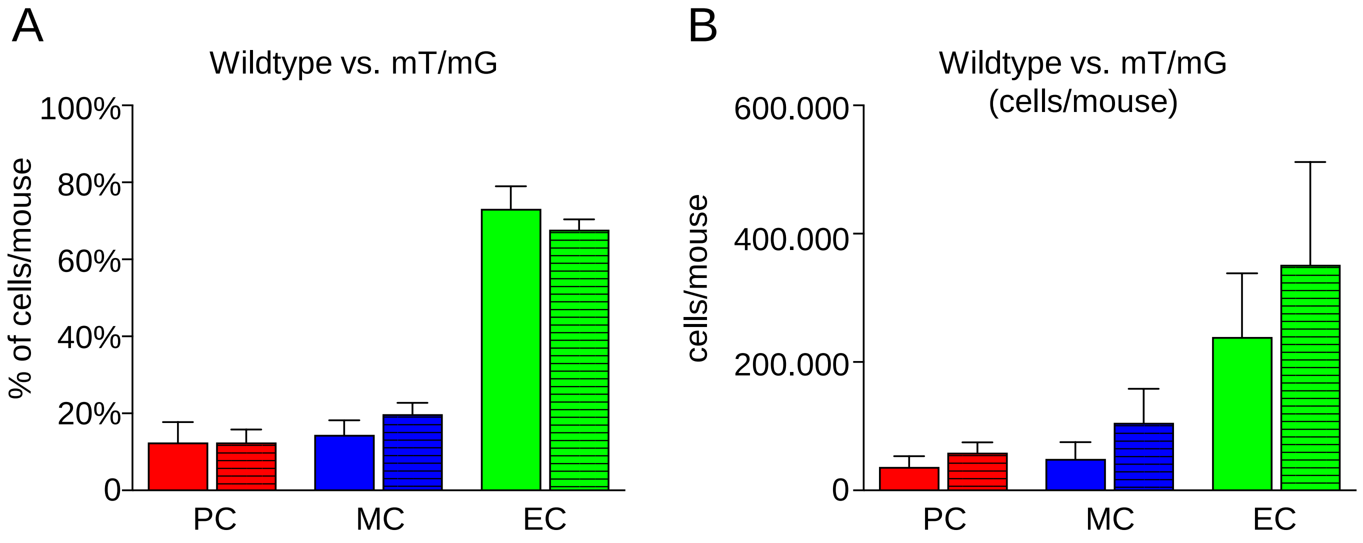


***Supplemental figure 3:*** **Comparison of FACS sort between mT/mG mice and wildtype mice**. (**A**) Percentage of podocytes, mesangial, and endothelial cells FACS-sorted from the isolated glomerular cells per mouse and (**B**) Absolute number of podocytes, mesangial, and endothelial cells FACS-sorted from the isolated glomerular cells per mouse, mean +/-SEM, n ≥ 23, pooled data from 18 independent experiments. Empty bars represent wildtype mice, stripped bars mT/mG mice.

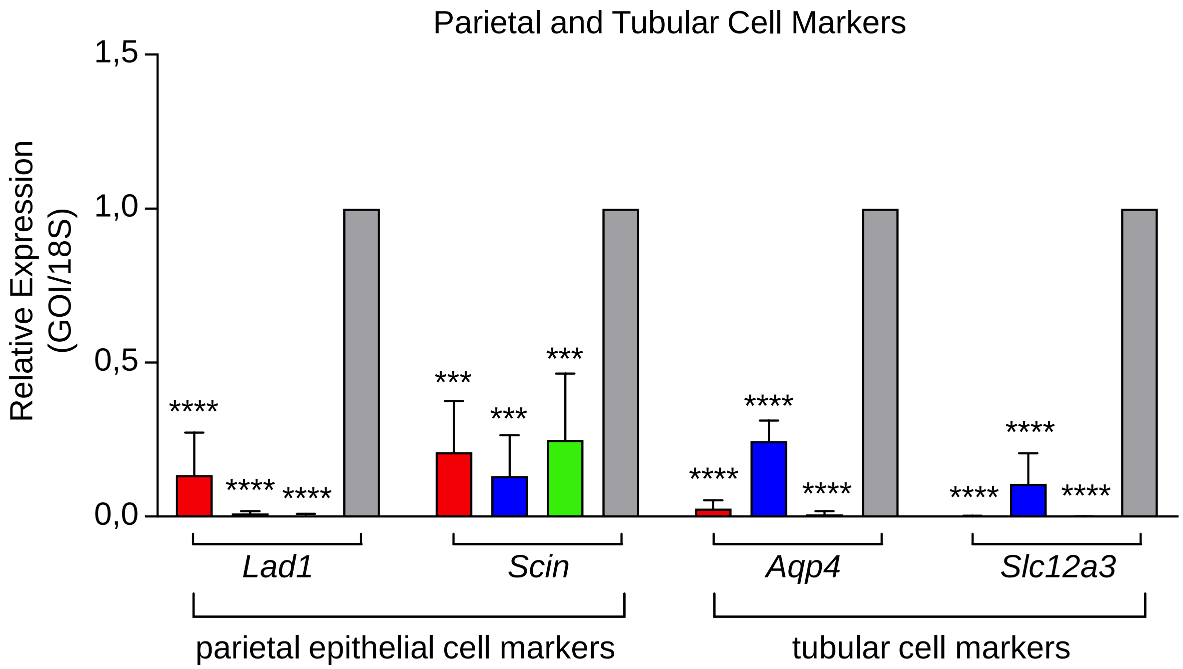


***Supplemental figure 4:*** **FACS sorted glomerular cells are not contaminated with parietal epithelial cells or tubular cells.** Real-time qPCR analysis exhibiting the expression of genes of interest (GOI) specific for parietal epithelial cells: *Lad1* (encodes for ladinin) and *Scin* (encodes for scinderin); and specific of tubular cells: *Aqp4* (encodes for aquaporin 4) and *Slc12a3* (encodes for sodium chloride channel NCC). mRNA was isolated from FACS-sorted podocytes (red bars), mesangial cells (blue bars), endothelial cells (green bars), and from isolated glomeruli (grey bars) derived from the same individual. ΔCT values were calculated using 18S as home keeper. ΔΔCT values were calculated as difference between glomerular and FACS-sorted cell type ΔCT. Displayed is the relative expression (RE, 2^(ΔΔCT)) of FACS-sorted cell types in percent of the glomerular RE. Note the significant decrease of cell-specific transcripts in the FACS sorted cell-types compared to the glomerulus, mean +/-SEM, ****p*<0.005; *****p*<0.0001, n=4, tested for statistical significance with ordinary one-way ANOVA and Bonferroni's multiple comparisons test.

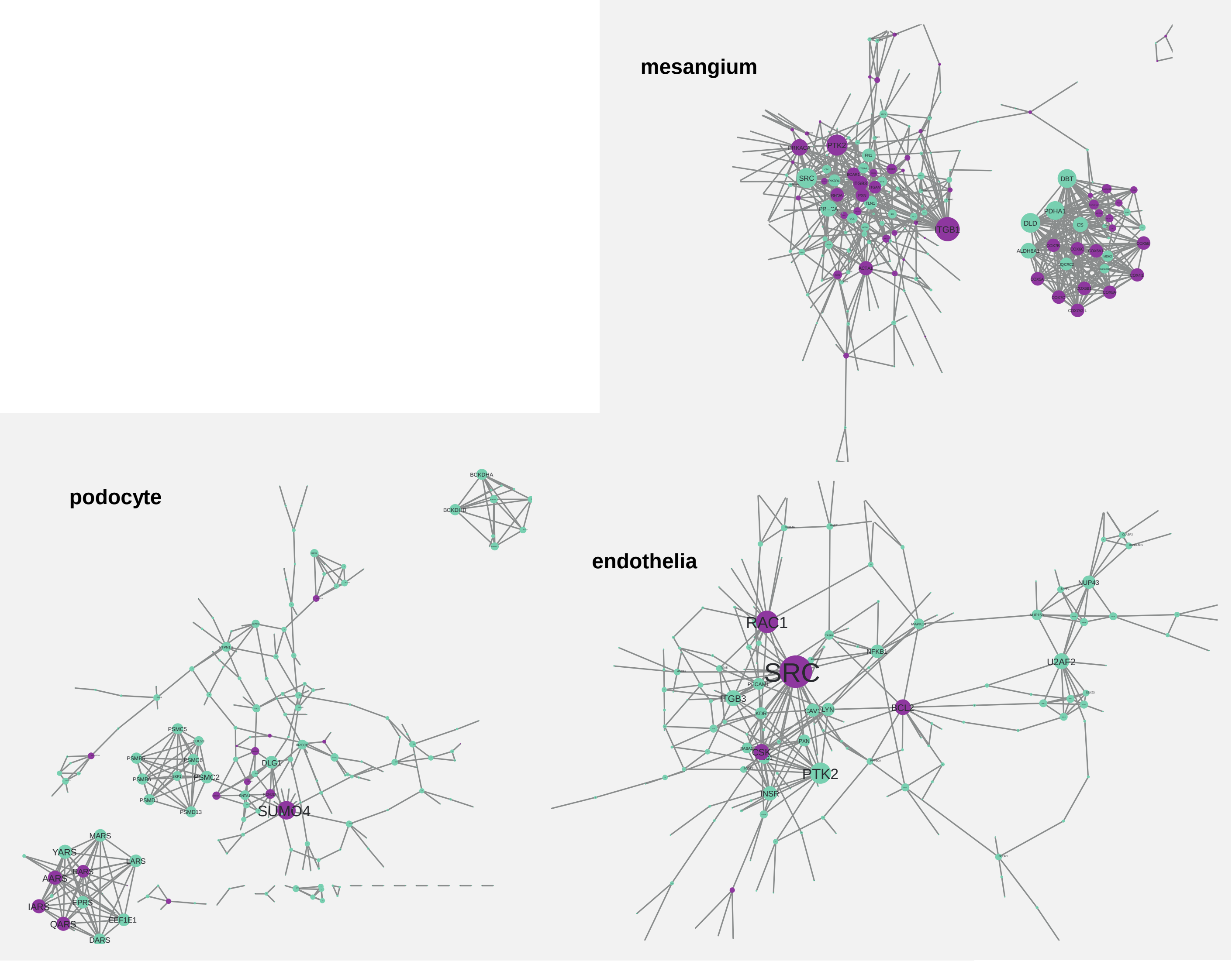


***Supplemental figure 5:*** **Tripartite cell type isolation and proteomic analysis from mT/mG mice.** Visualization of significant protein-protein interaction networks in original (green) and linker (violet) proteins.

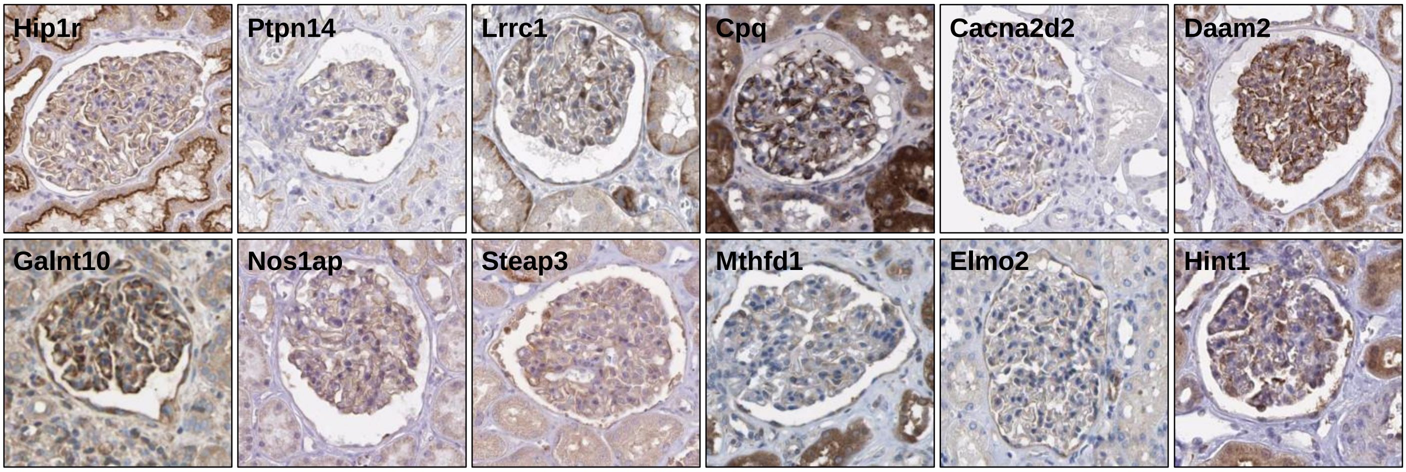


| Protein | Protein ID | Function (PMID) |
|---|---|---|
| **Hip1r** | Q9JKY5 | *Huntingtin-interacting protein 1-related protein*; Actin binding protein involved in clathrin mediated endocytosis (10613908); Isolog of huntingtin interacting protein (9852681) |
| **Ptpn14** | Q62130 | *Protein tyrosine phosphatase, nonreceptor-type 14*; Coimmunoprecipitates with VEGFR3 (136352) |
| **Lrrc1** | Q80VQ1 | *Leucine-rich repeat-containing protein 1;* Interacts with DLG1 (601014) and PSD95 (602887) |
| **Cpq** | Q9WVJ3 | *Carboxypeptidase Q*; Has significant glutamate carboxypeptidase activity (10206990) |
| **Cacna2d2** | Q6PHS9 | *Calcium channel, voltage-dependent, alpha 2/ delta subunit 2*; Confers modulation of presynaptic function (22678293) |
| **Daam2** | Q80U19 | *Dishevelled-associated activator of morphogenesis 2;* Member of the formin family, regulates actin and cytoskeletal dynamics and cell elongation (22275430) |
| **Galnt10** | Q6P9S7 | *Membrane-bound polypeptide N-acetylgalactosaminyltransferase;* Catalyzes the first step in mucin-type O-glycosylation of peptides in the Golgi (20977886) |
| **Flot2** | Q60634 | *Flotillin 2;* Human epidermal surface antigen-1, copurifies with Caveolin-1 (601047) |
| **Nos1ap** | Q9D3A8 | *Nitric oxide synthasae 1 (neuronal) adaptor protein*; Inhibition of L-type calcium channels (114205); Regulation of dendrite number (19553464) |
| **Steap3** | A0A0R4J1G9 | *Six-transmembrane epithelial antigen of prostate 3/Tsap6*; Involved in exosome secretion (18617898) |
| **Mthfd1** | Q922D8 | *Methylenetetrahydrofolate deyhdrogenase 1*; Mutations associated with combined immunodeficiency and megaloblastic anemia (27707659) and neuronal tube defects (12384833) |
| **Elmo2** | Q8BHL5 | *Engulfment and cell motility gene 2*; Phagocytosis and cell migration (11595183) |
| **Hint1** | P70349 | *Histidine triad nucleotide binding protein 1/PKCI1*; Interacts with PKC-beta, Mutations associated with axonal neuropathy and neuromyotonia (22961002) |

***Supplemental figure 6:*** **Identification of new podocyte enriched proteins.** To identify new podocyte enriched proteins, glomerular cell types proteome lists of individual mice were compared. For the comparisons we defined 1) a Student t-test difference cut-off of > 2 between podocytes and non-podocytes, 2) a negative search result in pubmed (https://www.ncbi.nlm.nih.gov/pubmed; i.e. with the keywords "protein of interest" and "podocyte") and 3) a validated podocyte expression pattern in the human protein atlas (https://www.proteinatlas.org/search), from which the histological micrographs are taken from. The table very briefly summarizes what is known for the identified proteins in other non-glomerular cell types, PMID = pub med identification number.

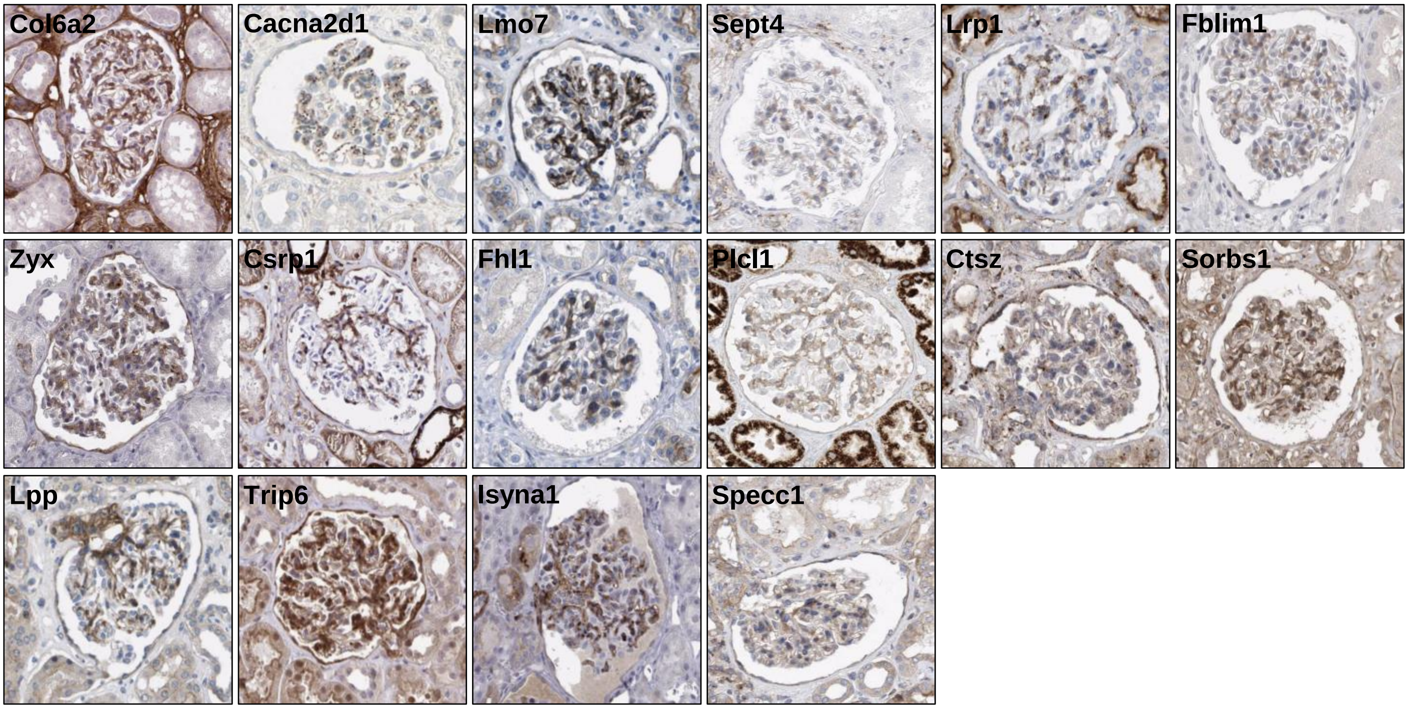


| Protein | Protein ID | Function (PMID) |
|---|---|---|
| **Col6a2** | Q02788 | *Collagen, type VI, alpha 2*; Integrity of muscle fibres (9817932); Mutations associated with muscle dystrophy (12840783) and myopathy (8782832) |
| **Cacna2d1** | O08532 | *Calcium channel, voltage-dependent, alpha 2/ delta subunit 1*; Alters properties of the pore-forming alpha 1 subunit of voltage gated calcium channels (10534405, 17088553) |
| **Lmo7** | F6VG99 | *Lim domain only 7*; Involved in protein-protein interactions, shuttles between cytoplasm and nucleus (17067998) |
| **Sept4** | P28661 | *Septin 4*; Polymerizing GTP-binding protine that scaffolds molecules beneath the plasma membrane; Colocalizes with alpha-synuclein in Lewy bodies (15737930; 17296554) |
| **Lrp1** | A0A0R4J0I9 | *Low density lipoprotein receptor related protein 1*; Interacts and traffics with APP (15749709) |
| **Fblim1** | Q71FD7 | *Filamin-binding LIM protein 1*; Localizes to cell-ECM adhesions, associates with actin filaments and is essential for cell shape modulation (12679033) |
| **Zyx** | Q62523 | *Zyxin*; Phosphoprotein concentrated at adhesion plaques and along actin filament bundles near where they insert at the adhesion plaques (8940160) |
| **Csrp1** | P97315 | *Cysteine- and glycine-rich protein 1*; Highly conserved, cell cycle regulated gene induced in the immediate early response to serum repletion in serum-starved noncycling cells (1385304) |
| **Fhl1** | A2AEX8 | *Four-and-a-half LIM domains 1*; Mutations associated with faciocardiomusculoskeletal syndrome (11102932), X-linked myopathy (18274675) |
| **Plcl1** | Q3USB7 | *Phospholipase C-like 1*; Negative regulation of bone formation (21757756) |
| **Ctsz** | Q9WUU7 | *Cathepsin Z*; member of papain family of cystein proteinases (9642240) |
| **Sorbs1** | Q62417 | *Sorbin and SH3-domains containing protein 1*; Required for insulin-stimulated GLUT4 translocation (11309621) |
| **Lpp** | Q8BFW7 | *LIM domain -containing preferred translocation partner in lipoma*; Most frequent translocation partner of HMGA2 in a subgroup of lipomas (15755872) |
| **Trip6** | Q9Z1Y4 | *Thyroid hormone receptor interactor 6*; Contains 2 LIM domains (601329) and is similar to zyxin (602002) |
| **Isyna1** | Q9JHU9 | *Inositol -3-phosphate synthase 1*; Component of plasma membrane phospholipids and functions as a cell signaling molecule |
| **Specc1** | A0A0J9YTU3 | *Sperm antigen with calponin homology and coiled-coil domains 1*; Function in head and neck tumors (22170762) |

***Supplemental figure 7:*** **Identification of new mesangial cell enriched proteins.** To identify new mesangial cell enriched proteins, glomerular cell types proteome lists of individual mice were compared. For the comparisons we defined 1) a Student t-test difference cut-off of > 2 between mesangial cell and non-mesangial cell, 2) a negative search result in pubmed (https://www.ncbi.nlm.nih.gov/pubmed; i.e. with the keywords “protein of interest” and “mesangial”) and 3) a validated mesangial expression pattern in the human protein atlas (https://www.proteinatlas.org/search), from which the histological micrographs are taken from. The table very briefly summarizes what is known for the identified proteins in other non-glomerular cell types, PMID = pub med identification number.

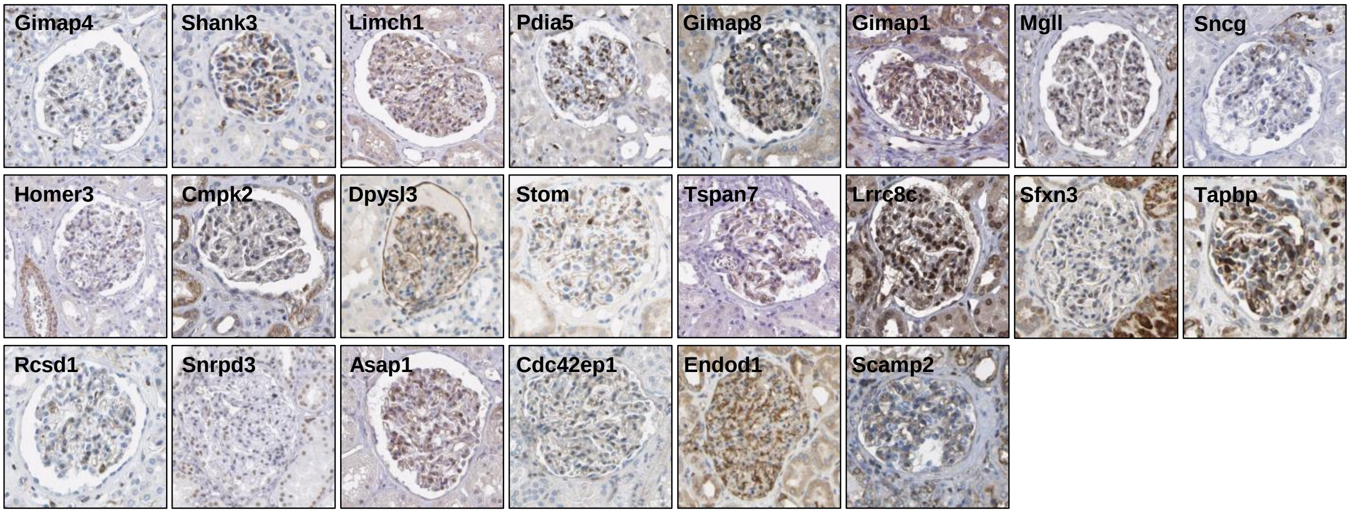


| Protein | Protein ID | Function (PMID) |
|---|---|---|
| **Gimap4** | Q99JY3 | *GTPase, Immunity-associated protein 4*; IFNg secretion regulation from T cells (25287446), regulation of T cell survival (16569770) |
| **Shank3** | Q4ACU6 | *SH3 and multiple ankyrin repeat domains 3*; Scaffolding protein enriched in postsynaptic densities of excitory synapses (26966193) |
| **Limch1** | Q3UH68 | *LIM and calponin homology domains-containing protein 1*; Actin stress fibre-associated protein that regulates nonmuscle myosin II (MYH9) activity (28338547) |
| **Pdia5** | Q921X9 | *Protein disulfide isomerase family A, member 5*; ER-resident protein that catalyzes oxidation, reduction, and isomerization of protein disulfide bonds which accelerates protein folding (24636989) |
| **Gimap8** | Q75N62 | *GTPase, Immunity-associated protein 8*; Regulator of lymphocyte survival and homeostasis (23454188) |
| **Gimap1** | Q3TFI6 | *GTPase, Immunity-associated protein 1;* Survival of peripheral T cells (287792288) and B cells (26621859) |
| **Mgll** | E9Q3B9 | *Monoglyceride lipase*; Functions together with hormone sensitive lipase to hydrolyze intracellular triglycerides stores to fatty acids and glycerol (11470505) |
| **Sncg** | Q9Z0F7 | *Synuclei gamma;* Highly expressed in neurons, correlation of overexpression and breastcancer progression (10813729) |
| **Homer3** | Q99JP6 | *Homer scaffold protein 3*; Cytoplasmic scaffolding protein that binds to metabotropic glutamate receptors (9069287) and interacts with Shank (19345194) |
| **Cmpk2** | Q3U5Q7 | *Cytidine monophosphate kinase 2*; Mitochondrial UMP-CMP kinase component of the salvage pathway for nucleotide synthesis (17999954) |
| **Dpysl3** | E9PWE8 | *Dihydropyrimidinase-like 3*; Role in neurite and axonal outgrowth (23568759), modulates mitosis migration and epithelial to mesenchymal transition (30498031) |
| **Stom** | P54116 | *Stomatin*; Modulates activity of anion exchanger 1(28387307), tumor suppressor (31949395) and enhancer of cell fusion when associated with lipid rafts (27663861) |
| **Tspan7** | Q3UHG5 | *Tetraspanin 7*; Contributes to molecular complexes that include b1-integrin (10655063), TSPN7 autoantibodies in diabetes type 1 (27221092) |
| **Lrrc8c** | Q8R502 | *Leucine-rich repeat-containing protein 8C*; B cell development (15094057) and adipocyte differentiation (15184384) |
| **Sfxn3** | Q91V61 | *Sideroflexin 3*; Mitochondrial serine transporter (30442778), a-synuclein dependent that regulates synaptic morphology (28049716) |
| **Tapbp** | Q3TCU5 | *Tap-binding protein*; Type 1 transmembrane protein encoded by an MHC-linked gene (8769474) in the ER, edits peptide loading onto MHC1 (32167472) |
| **Rcsd1** | Q3UZA1 | *RCSD domain-containing protein 1*; Interacts with F-actin capping protein Capz (601580) |
| **Snrpd3** | P62320 | *Small nuclear ribonucleoprotein polypeptide D3*; Part of snRNP core |
| **Asap1** | H3BL41 | *ARF GTPase-activating protein with SH3 domain, ankyrin repeat and PH domain 1*; Coordinates actin and membrane remodeling (25774636) |
| **Cdc42ep1** | Q91W92 | *CDC42 effector protein 1*; Mediates actin cytoskeletal reorganization at the plasma membrane (10430899) |
| **Endod1** | Q8C522 | *Endonuclease domain-containing 1*; Tumor suppressor in prostate cancer (28532481) |
| **Scamp2** | Q9ERN0 | *Secretory carrier membrane protein 2*; Regulates cell membrane targeting of NHE5 (19276089) and NKCC2 (21205824), role in granule exocytosis (12475951) |
| **Gimap4** | Q99JY3 | *GTPase, Immunity-associated protein 4*; IFNg secretion regulation from T cells (25287446), regulation of T cell survival (16569770) |

***Supplemental figure 8:*** **Identification of new glomerular endothelial cell enriched proteins.** To identify new glomerular endothelial cell enriched proteins, glomerular cell types proteome lists of individual mice were compared. For the comparisons we defined 1) a Student t-test difference cut-off of > 2 between endothelial and non-endothelial cell, 2) a negative search result in pubmed (https://www.ncbi.nlm.nih.gov/pubmed; i.e. with the keywords "protein of interest" and "endothelial") and 3) a validated glomerular endothelial expression pattern in the human protein atlas (https://www.proteinatlas.org/search), from which the histological micrographs are taken from. The table very briefly summarizes what is known for the identified proteins in other non-glomerular cell types, PMID = pub med identification number.

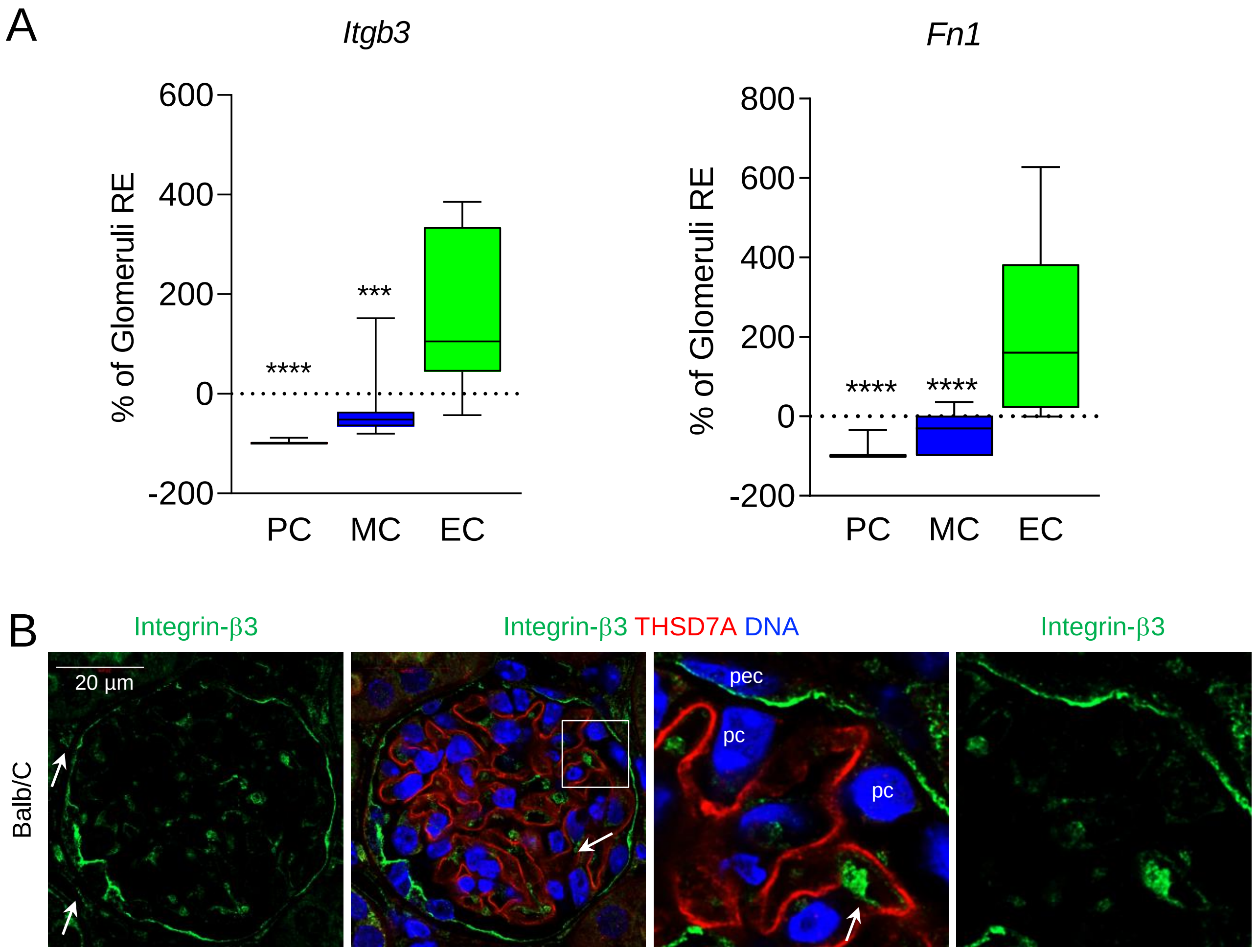


***Supplemental Figure 9:*** **Integrin-β3 and its ligand fibronectin 1 are predominantly expressed on glomerular endothelial cells.** (**A**) Integrin-β3 (*Itgb3*) and fibronectin (*Fn1*) transcript levels were quantified in FACS sorted glomerular cells relative to total glomeruli transcript levels in untreated BALB/c mice. Values are expressed as mean +/-SEM, N = 10 per group, ****p*<0.001 and *****p*<0.0001 to podocytes (PC), One Way ANOVA, Bonferroni's multiple comparison test. MC = mesangial cells, EC = glomerular endothelial cells. (**B**) Representative high-resolution confocal microscopy of integrin-β3 (green) expression in a BALB/c glomerulus in combination with THSD7A (red) to demarcate the podocyte foot processes and DNA (blue). Arrows point towards integrin-β3 expression in endothelial cells, pc = podocyte, pec = parietal epithelial cell.

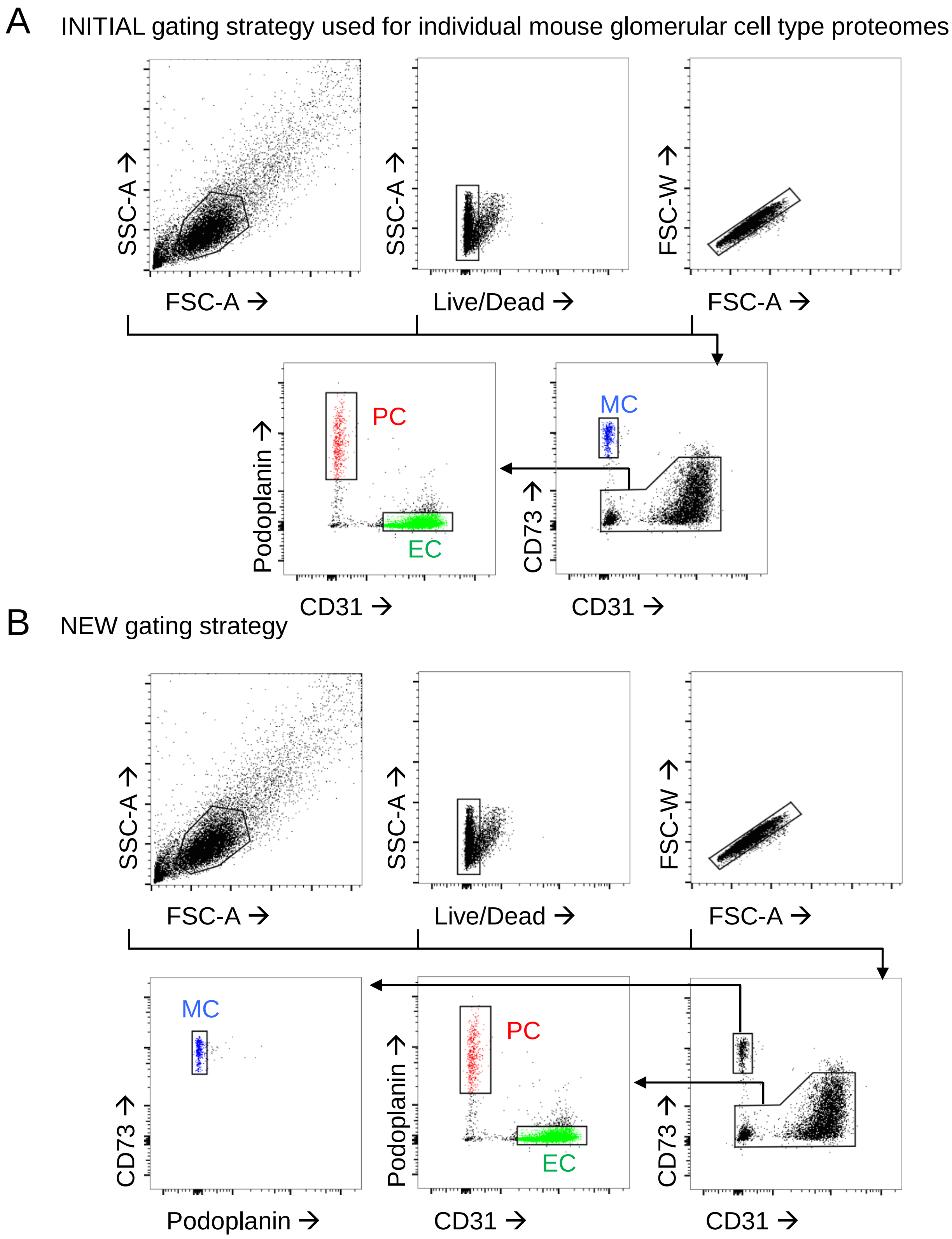


***Supplemental figure 10:*** **Comparison of old and new gating strategy. (A)** Gating strategy used for individual mouse proteome investigations in Figure 4 and (**B**) improved gating strategy removing podocyte contamination of mesangial cells used for all subsequent experiments.

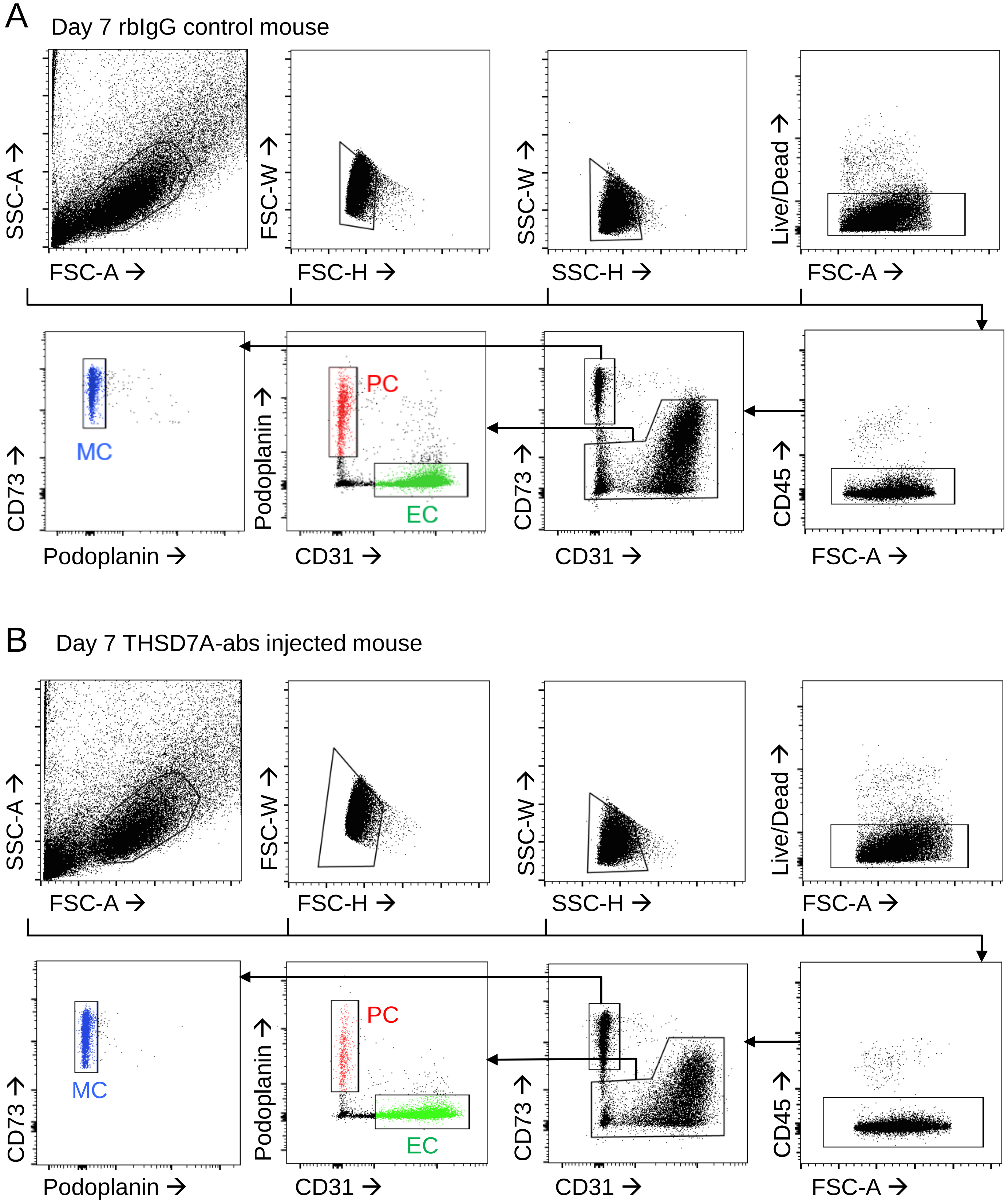


***Supplemental figure 11:*** **Gating strategy used for the separation of glomerular cell types in the mouse model of anti-THSD7A membranous nephropathy.** Individual mouse FACS plots are shown of a (**A**) unspecific rabbit IgG injected control mouse on day 7 and (**B**) a nephrotic anti-THSD7A-antibody injected mouse on day 7. CD45 was added to the FASC panel to remove potential contaminating leukocytes in the setting of glomerulonephritis.

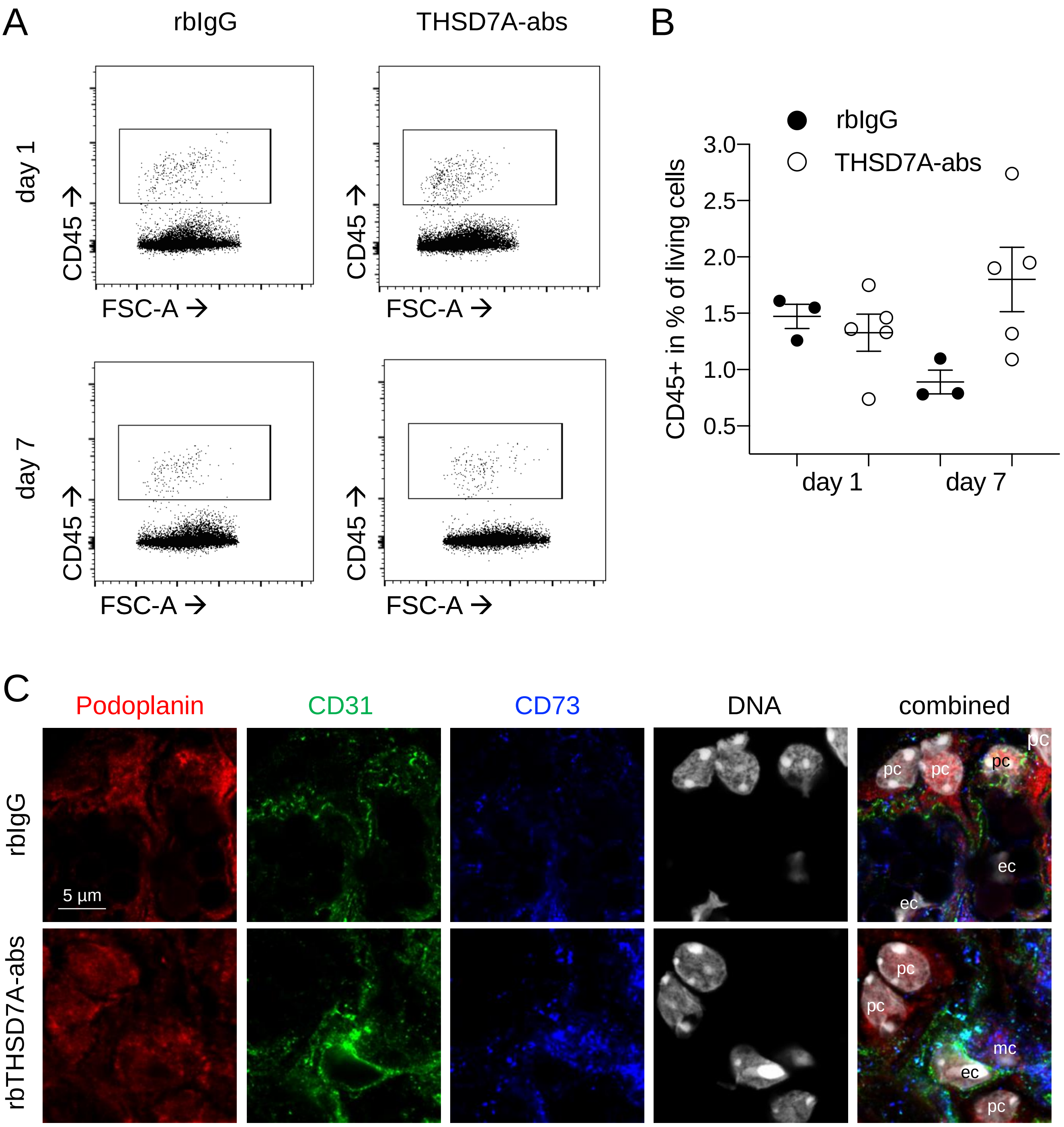


***Supplemental figure 12:*** **Leukocytes are not increased in glomeruli in the setting of experimental THSD7A-associated membranous nephropathy.** Experimental THSD7A membranous nephropathy was induced by injection of rabbit-anti-THSD7A antibodies or unspecific control rabbit IgG to male BALB/c mice and analyzed on day 1 and 7. (**A**) Representative FACS plots for CD45 at day 1 and day 7. (**B**) Quantification of CD45+ leukocytes in % of all living glomerular single cells on day 1 and day 7. (**C**) Confocal micrographs depicting labeled podocytes (podoplanin, red), endothelial cells (CD31, green) and mesangial cells (CD73, blue) in a 4% PFA fixed frozen section of a murine kidney, DNA was visualized using Hoechst, pc = podocyte, mc = mesangial cell, ec = endothelial cell.
